# GPR55 in B cells limits atherosclerosis development and regulates plasma cell maturation

**DOI:** 10.1101/2021.12.20.473518

**Authors:** Raquel Guillamat-Prats, Daniel Hering, Martina Rami, Carmen Härdtner, Donato Santovito, Petteri Rinne, Laura Bindila, Michael Hristov, Sabrina Pagano, Nicolas Vuilleumier, Sofie Schmid, Aleksandar Janjic, Wolfgang Enard, Christian Weber, Lars Maegdefessel, Alexander Faussner, Ingo Hilgendorf, Sabine Steffens

**Affiliations:** Institute for Cardiovascular Prevention (IPEK), Ludwig-Maximilians-Universität (LMU) Munich, Munich, Germany; German Center for Cardiovascular Research (DZHK), Partner Site Munich Heart Alliance (MHA), Munich, Germany; Department of Cardiology and Angiology I, Heart Center and Faculty of Medicine, University of Freiburg. Freiburg, Germany; Institute for Experimental Cardiovascular Medicine, Heart Center and Faculty of Medicine, University of Freiburg, Freiburg, Germany; Institute for Genetic and Biomedical Research, UoS of Milan, National Research Council, Milan, Italy; Institute of Biomedicine, University of Turku, Turku, Finland; Institute of Physiological Chemistry, University Medical Center of the Johannes Gutenberg University Mainz, Mainz, Germany; Division of Laboratory Medicine, Department of Genetics and Laboratory Medicine, Geneva University Hospitals, Geneva, Switzerland; Department of Vascular and Endovascular Surgery, Klinikum rechts der Isar - Technical University Munich (TUM), Munich, Germany; Anthropology and Human Genomics, Faculty of Biology, Ludwig-Maximilians University, Martinsried, Germany; Department of Biochemistry, Cardiovascular Research Institute Maastricht (CARIM), Maastricht University Medical Centre, 6229 ER Maastricht, The Netherlands; Munich Cluster for Systems Neurology (SyNergy), Munich, Germany

## Abstract

Identifying novel pathways regulating the adaptive immune response in chronic inflammatory diseases such as atherosclerosis is of particular interest in view of developing new therapeutic drugs. Here we report that the lipid receptor GPR55 is highly expressed by splenic B cells and inversely correlates with atheroma plaque size in mice. In human carotid endarterectomy specimen, GPR55 transcript levels were significantly lower in unstable compared to stable carotid plaques. To study the impact of GPR55 deficiency in atherosclerosis, we crossed *Gpr55* knockout mice with apolipoprotein E (*ApoE*) knockout mice and subjected the mice to Western diet for 4 to 16 weeks. Compared to *ApoE*^-/-^ controls, *ApoE^-/-^Gpr55^-/-^* mice developed larger plaques with increased necrotic core size, associated with elevated circulating and aortic leukocyte counts. Flow cytometry, immunofluorescence and RNA-sequencing analysis of splenic B cells in these mice revealed a hyperactivated B cell phenotype with disturbed plasma cell maturation and immunoglobulin (Ig)G antibody overproduction. The specific contribution of B cell GPR55 in atherosclerosis was further studied in mixed *Gpr55^-/-^*/*µMT* bone marrow chimeras on low density receptor deficiency (*Ldlr^-/-^*) background, revealing that B-cell specific depletion of *Gpr55* was sufficient to promote plaque development. Conversely, adoptive transfer of wildtype B cells into *ApoE^-/-^Gpr55^-/-^* mice blunted the proatherogenic phenotype. *In vitro* stimulation of splenocytes with the endogenous GPR55 ligand LPI promoted plasma cell proliferation and enhanced B cell activation marker expression, which was inhibited by the GPR55 antagonist CID16020046. Collectively, these discoveries provide new evidence for GPR55 as key modulator of the adaptive immune response in atherosclerosis. Targeting GPR55 could be useful to limit inflammation and plaque progression in patients suffering from atherosclerosis.

## Introduction

Cardiovascular diseases (CVDs) related to atherosclerosis are the leading cause of death worldwide. Atherosclerosis is a chronic and multifactorial disease involving both arms of our immune system, the innate and adaptive immunity.^1–3^ Recent advances in analytical approaches such as single cell sequencing and mass cytometry have deepened our knowledge about the cellular heterogeneity within the atherosclerotic lesions,^4^ supporting the relevance of B lymphocytes in the pathophysiology of the disease.

Diverse subset-specific functions of B cells in atherosclerosis have been described over the last years, reporting both pro- and antiatherogenic properties.^5–8^ For example, B1a cells are considered as main producers of naturally occurring IgM antibodies arising without antigen-mediated induction. These antibodies recognize oxidation-specific epitopes (OSE) that are present on lipoprotein particles as well as apoptotic cells, thereby limiting atherosclerotic plaque progression.^8^ The B2 lineage can be separated into marginal zone (MZ) and follicular (FO) cells. While MZ cells are positioned at the splenic red pulp border to scan the blood for circulating antigens, FO cells interact with T follicular helper (TFh) cells to initiate the germinal center (GC) response.^9^ This culminates in the final stage of B cell maturation, which involves the rapid proliferation of plasmablasts secreting low antibody levels, somatic hypermutations in the B cell receptor (BCR) region and selection of high antigen affinity clones to ultimately differentiate into long-lived plasma cells.^9^ Plasma cells have a high protein synthesis capacity and secrete large amounts of antibodies into the circulation, which depends on a set of transcription factors, in particular PR domain zinc finger protein 1 (BLIMP1), interferon regulatory factor 4 (IRF4) and X-box binding protein 1 (XBP1). Antigen-specific antibodies of various isotypes, including IgGs, are produced by class-switching of the high affinity B cell clones.

In the context of atherosclerosis, IgGs have been reported to form immune complexes with oxLDL and promote macrophage inflammatory responses.^10^ On the other hand, high-throughput single-cell analysis of the atherosclerosis-associated antibody repertoire recently discovered atheroprotective anti-ALDH4A1 autoantibodies directed against a mitochondrial dehydrogenase.^11^ Moreover, MZ B cells have been shown to negatively control proatherogenic Tfh responses to hypercholesterolemia,^12^ which underlines the complexity of B cell subset-specific functions and autoantibodies in atherosclerosis.

B cell activation involves antigen binding to the BCRs, which are membrane-bound immunoglobulins with unique epitope-binding sites expressed by each clone. The BCRs form a signaling complex with other receptors on the B cell surface, including among other, receptor-type tyrosine-protein phosphatase C (PTPRC, also known as B220).^10^ Under hypercholesterolemic conditions, B cells may become hyperactivated with deregulation of plasma cell formation, thereby promoting atherosclerosis.^10, 13^

GPR55 is a G protein-coupled orphan receptor expressed by various leukocyte subsets, including B and T cells, but also natural killer cells, monocytes, macrophages, and neutrophils. According to murine immune cell transcriptomic data (Immgen.org),^14^ GPR55 is highly expressed by splenic plasma cells (PC) and, to a lower extent, MZ B cells. The most potent endogenous GPR55 ligand identified so far is lysophosphatidylinositol (LPI).^15–17^ Immunological functions for GPR55 have been described, amongst others, in γδT cells residing in intestinal lymphoid organs, where *Gpr55* deficiency or short-term antagonist treatment protected mice from nonsteroidal anti-inflammatory drug-induced increases in intestinal permeability.^18^

In THP1-derived macrophages, GPR55 stimulation promoted the accumulation of oxidized LDL and blocked cholesterol efflux, while GPR55 antagonism counteracted these effects.^19^ Conversely, in aortic endothelial cells, antagonizing GPR55 with CID16020046 prevented ox-LDL-induced inflammatory stress.^20^ *In vivo* treatment of atherosclerosis-prone *ApoE^-/-^* mice with CID16020046 reduced neutrophil recruitment and plaque infiltration, while no effects on plaque size were detectable.^21^ However, the role of GPR55 in regulating adaptive B cell responses in the context of atherosclerosis has not been investigated so far.

Using a global *Gpr55* knockout mouse model, B cell specific deletion and gain of function experiments, we report here for the first time that GPR55 plays a role in regulating B cell activation and maturation during hypercholesterolemia, which crucially affects atherosclerotic plaque development. Our findings provide further insight into the complex regulation of humoral immunity and might be of broader relevance beyond atherosclerosis, such as autoimmune disorders.

## Results

### Modulation and role of the GPR55-LPI axis in atherosclerosis

To address the regulation of the GPR55-LPI axis during hypercholesterolemia, we fed *ApoE^-/-^* mice with Western diet (WD) for 4 or 16 weeks, respectively, and measured LPI plasma levels by liquid chromatography-tandem mass spectrometry. Circulating LPI was elevated during the early stage of atherogenesis and returned to baseline concentrations after 16 weeks WD (Figure 1A). Given that GPR55 is highly expressed by splenic B cells, we focused on the spleen to assess a modulation of the receptor and the LPI synthesizing enzyme phospholipase DDHD domain containing 1 (DDHD1) during atherosclerosis onset and progression. In accordance with the modulation of plasma LPI levels, *Ddhd1* was upregulated after 4, but not 16 weeks WD (Figure 1B). A similar pattern for the splenic *Gpr55* mRNA expression was observed (Figure 1C). At the 4-week time point, the splenic GPR55 expression inversely correlated with aortic root atherosclerotic plaque size (Figure 1D), suggesting that GPR55 signaling may exert an early protective effect on plaque development.

**Figure 1:**
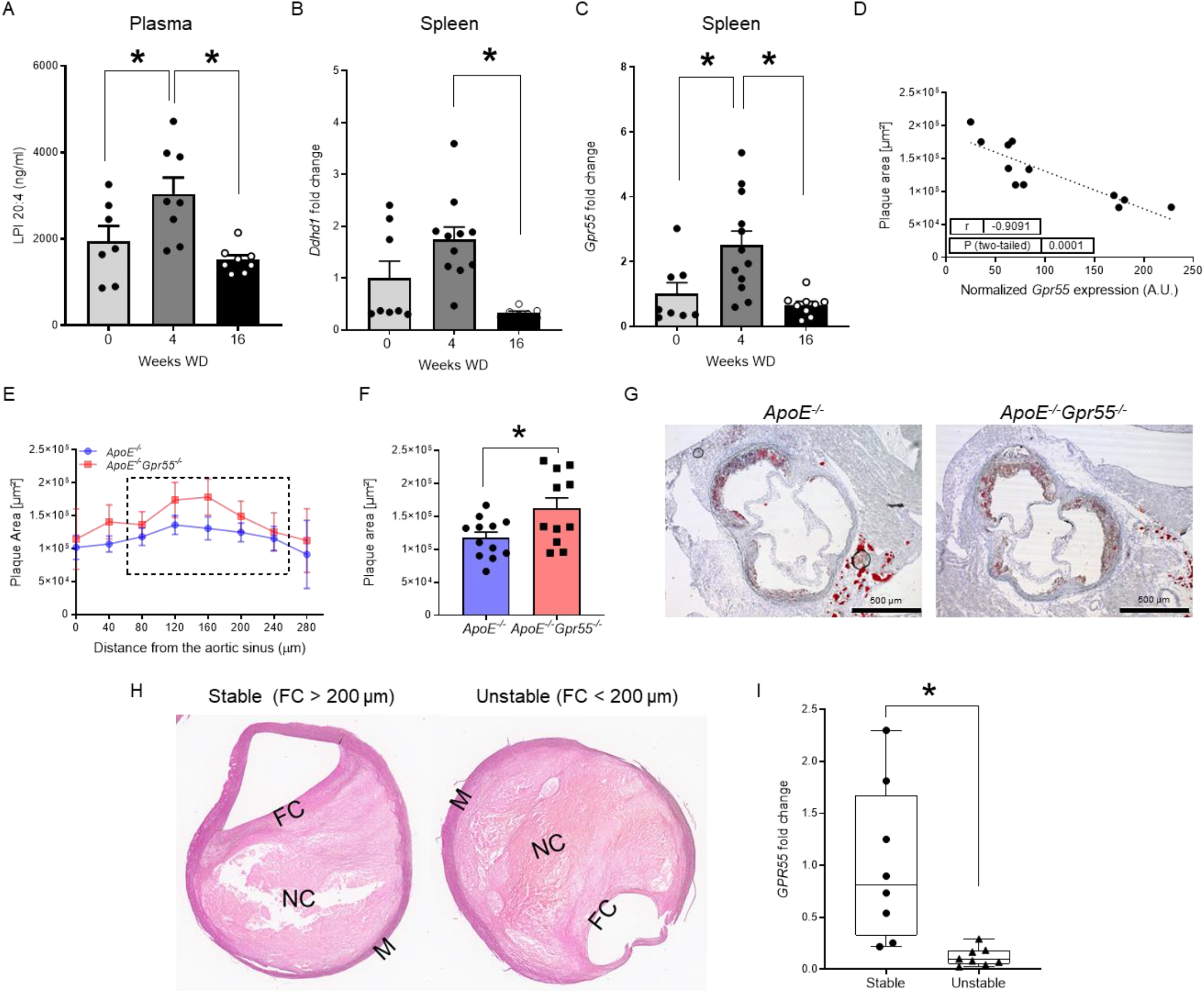
Regulation and function of GPR55 signaling in atherosclerosis. (**A-D**) Plasma, spleens and aortic roots were collected from *ApoE^-/-^* mice at baseline or after 4 and 16 weeks WD to determine (**A**) lysophosphatidylinositol (LPI) plasma concentrations or (**B-C**) relative splenic mRNA expression of the gene encoding the LPI-synthesizing enzyme DDHD1 and the LPI receptor GPR55. (**D**) Splenic *Gpr55* mRNA expression values were correlated to the aortic root plaque areas of the same mice. (**E**) Plaque area per aortic root section of female *ApoE^-/-^* and *ApoE^-/-^Gpr55^-/-^* mice after 4 weeks WD (n=12 per group). The dotted square indicates the sections used for calculating the average plaque area per animal shown in (**F**). (**G**) Representative Oil-Red-O stains of aortic roots after 4 weeks WD. (**H**) Representative pictures of human stable and unstable plaques (FC=fibrous cap, NC=necrotic core, M=media) obtained from the Munich Vascular Biobank. (**I**) Human *GPR55* mRNA expression evaluated by qPCR in stable vs. unstable/ruptured carotid artery plaque corrected by *RPLPO* used as housekeeping control. Each dot represents one patient from the same Biobank. Mouse data shown in A-F were combined from 3 independent experiments (mean ± SEM; each point represents one animal). Unpaired Student’s t-test was used to determine the significant differences *p < 0.05 vs the indicated group.

The causal implication of GPR55 in atherogenesis was substantiated in *ApoE^-/-^Gpr55^-/-^* mice receiving 4 weeks WD, which developed larger plaques within aortic roots compared to corresponding *ApoE^-/-^* controls (Figure 1E-G). These findings suggest that GPR55-signaling counterbalances plaque development, at least in early disease stage. After 16 weeks WD, a difference in aortic lesion area was no longer observed (Supplementary Figure 1A-B). In the descending aorta, however, the plaque burden was still higher in *ApoE^-/-^Gpr55^-/-^* mice (Supplementary Figure 1C). This might reflect stage dependent effects of GPR55 during atherosclerosis, since plaque development in descending aortas is generally less advanced compared to aortic root plaques in mouse models of atherosclerosis.^22^ In support of this hypothesis, 16-weeks-plaques of *Gpr55*-deficient mice exhibited an increased necrotic core and collagen area, but decreased relative plaque content of macrophages and lipids, indicative of a more advanced plaque phenotype (Supplementary Figure 1D to G).

To validate the expression and possible modulation of GPR55 in human plaques, we analyzed the transcript levels in stable and unstable carotid artery plaques collected from patients undergoing endarterectomy. The patient’s characteristics of both groups are summarized in Supplementary Table 1. The plaque phenotype was assigned based on the morphology following the American Heart Association (AHA) classification^23^ with or without a vulnerable fibrous cap according to Redgrave et al.^24^ (Figure 1H and Supplementary Table 3 with the patient characteristics). The qPCR analysis revealed reduced *GPR55* expression in unstable compared to stable plaques (Figure 1I), which might suggest that high expression levels of this receptor may counterbalance the patient’s risk for developing an acute cardiovascular event.

### B cell *Gpr55* expression and functionality

We next assessed the expression profile of *Gpr55* in sorted murine blood leukocytes of *ApoE^-/-^* mice by digital droplet PCR (ddPCR) and qPCR. The rank order of *Gpr55* expression in the different populations was B cells > T cells > neutrophils > monocytes, which was consistently confirmed by both techniques (Figure 2A). By *in situ* hybridization, we localized *Gpr55* expression in the splenic B cell FO and GC areas (Figure 2B).

**Figure 2:**
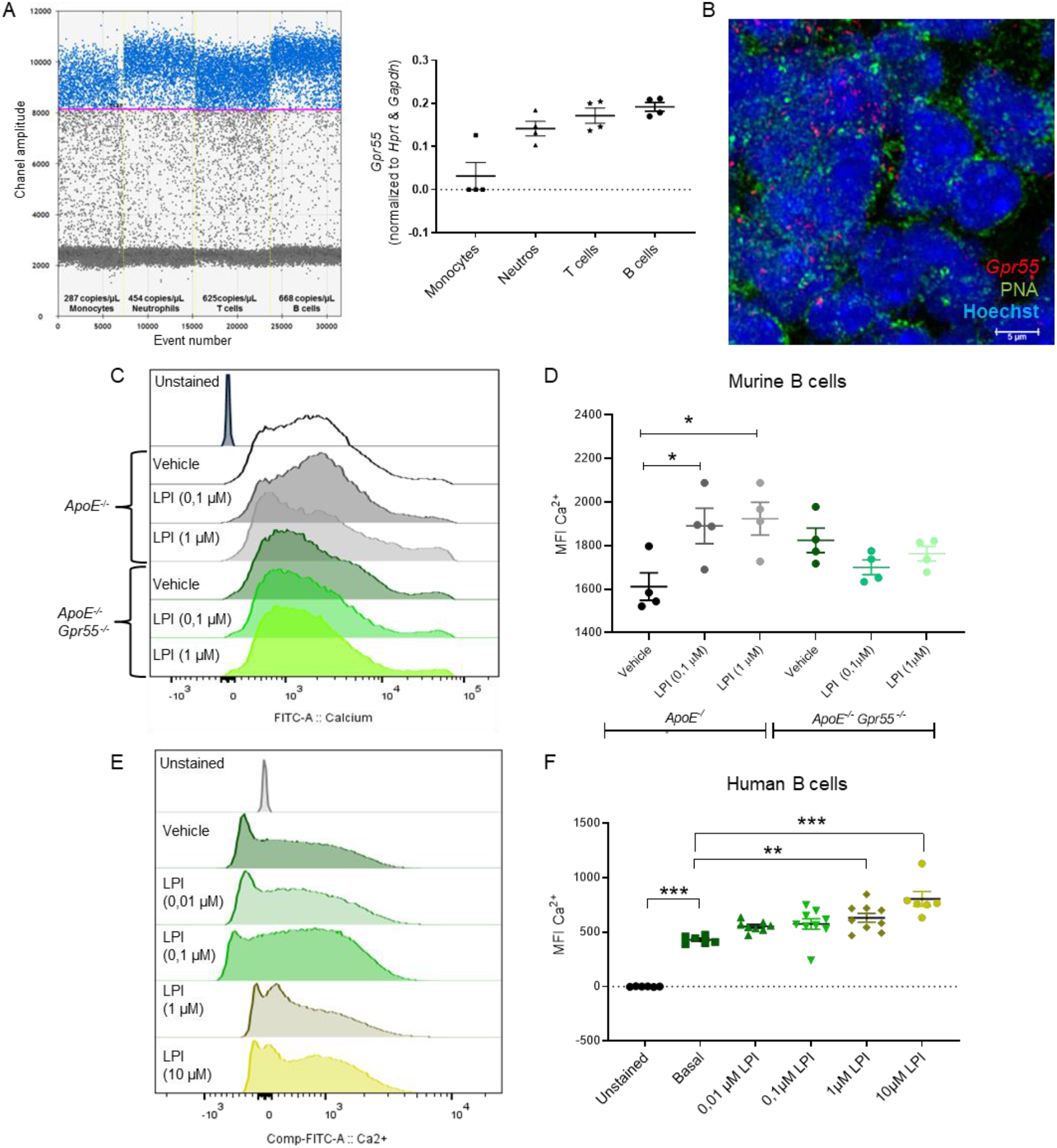
GPR55 expression and GPR55-dependent Ca^2+^ responses in murine and human B cells. (**A**) *Gpr55* mRNA expression determined by digital droplet PCR (left) and qPCR (right) in sorted circulating leukocyte subsets. (**B**) *In situ* hybridization to detect *Gpr55* (red) in splenic GC areas (PNA^+^, green; nuclei, blue). (**C**) Representative histogram and (**D**) average mean fluorescence intensity (MFI) of intracellular Ca^2+^ sensor for 30 seconds in circulating B cells of *ApoE^-^*^/-^ and *ApoE^-^*^/-^*Gpr55^-/-.^*mice in response to LPI stimulation. (**E**) Representative histogram and (**F**) average MFI of intracellular Ca^2+^ sensor in human B cells at baseline conditions and after addition of vehicle or LPI. Data were combined from 2 independent experiments (mean ± SEM; each point represents one mouse or human donor, respectively). One-Way-ANOVA followed by a post-hoc Newman–Keuls multiple-comparison test was used to evaluate the significant differences *p < 0.05, ** p < 0.01, *** p < 0.001 vs the indicated group.

The functional GPR55 signaling response in splenic B cells was subsequently validated by measuring intracellular Calcium (Ca^2+^) increases in response to stimulation with the endogenous GPR55 ligand LPI. Dose-dependent increases in intracellular Ca^2+^ were observed in *ApoE^-/-^,* but not in *ApoE^-/-^Gpr55^-/-^* mice (0,1 µM and 1 µM; Figure 2C-D). Likewise, human peripheral blood B cells showed a dose-dependent Ca^2+^ response to LPI, indicating a functional LPI-GPR55 signaling pathway in human B cells (1 µM and 10 µM; Figure 2E-F).

### Global *Gpr55* deficiency leads to increased metabolic dysfunction and inflammation during hypercholesterolemia

Global *Gpr55* deficiency was associated with pronounced metabolic changes, in particular increased body weight compared to aged-matched *ApoE*^-/-^ animals, which was already notable at baseline and consistently evident after 4 or 16 weeks WD (Supplementary Figure 2A-C). Although plasma cholesterol levels were comparable to *ApoE^-/-^* controls at all time points (Supplementary Figure 2D), the liver total cholesterol was slightly increased after 4 weeks WD (Supplementary Figure 2E). The hepatic qPCR gene expression analysis after 4 weeks WD revealed a significant transcriptional upregulation of several metabolic markers, indicating enhanced lipid synthesis, uptake and efflux in *Gpr55*-deficient mice (Supplementary Figure 2F). Furthermore, the Oil Red O staining analysis of liver sections revealed an enhanced lipid droplet accumulation after 16 weeks WD (Supplementary Figure 2G-H). These findings are in agreement with recent reports on pro-steatotic effects of GPR55.^25^

We next investigated the effect of global GPR55 deficiency on vascular and systemic inflammation under conditions of hypercholesterolemia. Compared to *ApoE^-/-^* controls, *ApoE^-/-^Gpr55^-/-^* mice had a hyper-inflammatory phenotype with an increase in all major circulating and aortic leukocyte subsets (Supplementary Figure 2I-K); accompanied by an amplified aortic gene expression profile of key pro-inflammatory markers (Supplementary Figure 2L).

Similar effects on plaque development and metabolic changes, as detailed in female mice, were observed in male *Gpr55*-deficient mice (Supplementary Figure 3A-C).

### *Gpr55* deficiency is associated with enhanced antibody responses during hypercholesterolemia

To seek for a possible underlying reason for the excessive pro-inflammatory phenotype and considering that GPR55 is mostly expressed by splenic B cells (Figure 2A), we performed a detailed flow cytometric profiling of B cell populations. Since splenic GPR55 expression was upregulated after 4 weeks WD, but not at the later stage (Figure 1C), we subsequently focused on this time point for an in-depth profiling of the *Gpr55*-deficient B cell phenotype. The flow cytometric analysis showed a diminished number of splenic B1a counts in *ApoE*^-/-^*Gpr55*^-/-^ mice compared to *ApoE*^-/-^ controls, while B2 marginal zone (MZ) cell counts were increased (Figure 3A). The most pronounced changes were reductions of splenic germinal center (GC) B cell and PC counts in *Gpr55*-deficient mice (Figure 3A). A similar pattern was observed when comparing the relative frequencies among splenic B cell subsets (Figure 3B-C). B1 cells in *ApoE^-/-^Gpr55^-/-^* mice were shifted to a decreased proportion of the anti-atherogenic B1a subset with increased B1b subset (Figure 3B-C). Furthermore, there was a decreased proportion of the follicular (FO) subset and increased proportion of the proatherogenic MZ subset (Figure 3D-E). These flow cytometric data were supported by immunohistological analysis of splenic tissue sections, indicating that B cell follicular and GC areas were less well-defined in mice lacking a functional GPR55 receptor (Figure 3F).

**Figure 3:**
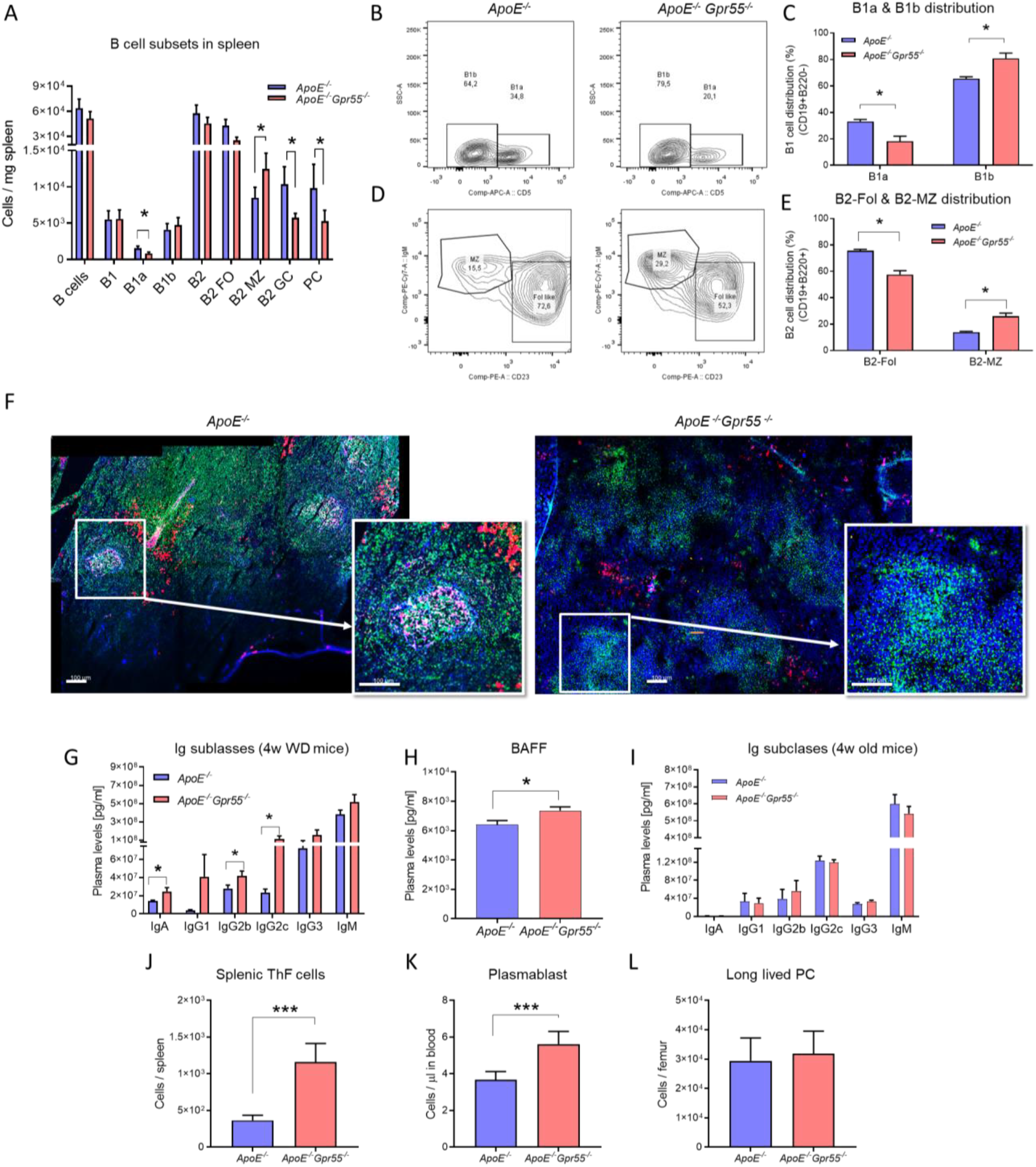
B cell immunoprofiling of global *Gpr55*-deficient mice after 4 weeks WD. (**A**) Splenic B cell subsets quantified by flow cytometry. (**B, C**) Representative gating strategy and relative proportion of B1 cell subsets. (**D, E**) Representative gating strategy and relative proportion of B2 cell subsets. (**F**) Confocal imaging of spleen sections stained for B cell markers PNA (GC, green), IgM (MZ or memory B cells, red), CD23 (FO B cells, pink) and nuclei (blue). (**G**) Plasma Ig titers and (**H**) B cell activating factor (BAAF) concentrations after 4 weeks WD. (**I**) Plasma Ig titers in young 4-week-old mice. (**J**) Number of splenic T follicular helper (Tfh) cells, (**K**) circulating plasmablasts and (**L**) long lived plasma cells (LLPC) in bone marrow assessed by flow cytometry after 4 weeks WD. Data were combined from 2 independent experiments (mean ± SEM; A-G, n=15-16; H-L, n=6-8 per group). Unpaired Student’s t-test was used to determine the significant differences *p < 0.05 and ***p < 0.001 vs *ApoE^-/-^*.

Despite the reduced PC counts, *Gpr55*-deficient mice had elevated titers of IgA antibodies and several IgG subclasses (Fig. 3G). The most pronounced increase was detected in IgG2c antibody titers. Comparable increases of IgG subclass titers were observed in male *Gpr55*-deficient mice (Supplementary Figure 3D). Moreover, *Gpr55*-deficient mice had elevated plasma levels of B cell activating factor (BAFF; Figure 3H), which is crucial for B cell survival, altogether indicating a hyperactivated B cell response during atherosclerosis in the absence of GPR55 signaling. To clarify whether *Gpr55* deficiency per se affected B cell maturation and antibody secretion, independent of antigens such as OSE arising during hypercholesterolemia, we also quantified antibody titers in the plasma of young mice at 4 weeks of age. No differences in antibody titers between young *ApoE*^-/-^ and *ApoE*^-/-^*Gpr55*^-/-^ were detectable, suggesting that proatherogenic conditions trigger the hyperactivated phenotype of B cells when functional GPR55 signaling is lacking (Figure 3I).

The hyperactivated B cell state was linked to an increased number of splenic follicular T helper (Tfh) cells and circulating plasmablasts in *Gpr55*-deficient mice (Figure 3 J and K), while long-lived PC counts in the bone marrow were unchanged (Figure 3L). Taken together, these data suggest an unbalanced humoral response in atherosclerotic mice lacking functional GPR55 signaling, possibly due to disturbed checkpoints controlling the B cell differentiation from a highly proliferative into a specialized antibody secreting cell.

### Bulk RNA-sequencing analysis of splenic B cells

To understand in more detail the role of GPR55 signaling in splenic B cells, we sorted splenic CD19^+^ B cells from 6 *ApoE^-/-^* and 6 *ApoE^-/-^Gpr55^-/-^* mice after 4 weeks of WD and performed prime-seq, a bulk RNA-sequencing protocol.^26^ We found 460 differentially expressed genes (DEGs) between *ApoE^-/-^* and *ApoE^-/-^Gpr55^-/-^* mice that were enriched in B and T cell activation, cellular response to stress and intracellular signal transduction (Figure 4A and Supplementary Figure 4A-B). The main DEGs associated to each GO pathway were validated by qPCR, confirming a decrease of *Fcer2a*/CD23 and *Ptprc*/LPAP, together with changes in *Zbtb20*, *Xbp1* and *Bcl6*, which are important transcription factors involved in B cell maturation and PC differentiation^10^ (Figure 4B). To study in more depth the transcriptomic changes linked to altered PC maturation in *Gpr55*-deficient mice, we also performed bulk RNA-sequencing of sorted splenic PC, blood plasmablasts and bone marrow long-lived PC from *ApoE^-/-^* versus *ApoE^-/-^Gpr55^-/-^* mice. The integration of differentially regulated GO pathways in a chord diagram confirmed that the three populations show overlapping regulated pathways such as B and T cell activation, disrupted Ig production and activated cell stress genes, confirming that B cell maturation and PC function is generally affected by G*pr55* deficiency (Figure 4C). In *ApoE^-/-^* controls, the total splenic G*pr55* expression directly correlated with *Fcer2a*/CD23 and *Ptcarp/*LPAP (Figure 4D-E). B2-FO cells downregulate *Fcer2a*/CD23 expression on the cell surface when they go into memory B cell differentiation.^27^ *Ptprc*/LPAP expression was lower in *Gpr55*^-/-^ mice, which is in line with previous data reporting that LPAP deficiency is associated with increased MZ B2 cell numbers.^28^ Evaluation of transcriptional factors involved in B cell biology revealed lower expression of *Zbtb20* in spleens of *Gpr55^-/-^* mice, while *Xbp1* expression was 2-fold increased. *Zbtb20* is a key player in late B cell differentiation and is highly expressed by GC and PC.^29^ The decreased levels in *Gpr55*-deficient mice are in line with the lower numbers of GC B cells and disturbed PC differentiation (Figure 2C).^29^ Overexpression of *Xbp1* promotes long-lived PC differentiation and is crucial for switching into secretory cells, releasing large quantities of antibodies, inducing endoplasmic reticulum (ER) remodeling, autophagic pathways, and the induction of the unfolded protein response (UPR).^30^ To substantiate these findings, we compared the CD23 protein expression on splenic B cells, confirming reduced surface expression in *Gpr55*-deficient mice (Figure 4G). *In vitro* stimulation of splenocytes with the GPR55 agonist LPI upregulated CD23 surface expression levels in B cells, with a maximum effect observed after 30 min (Figure 4F).

**Figure 4:**
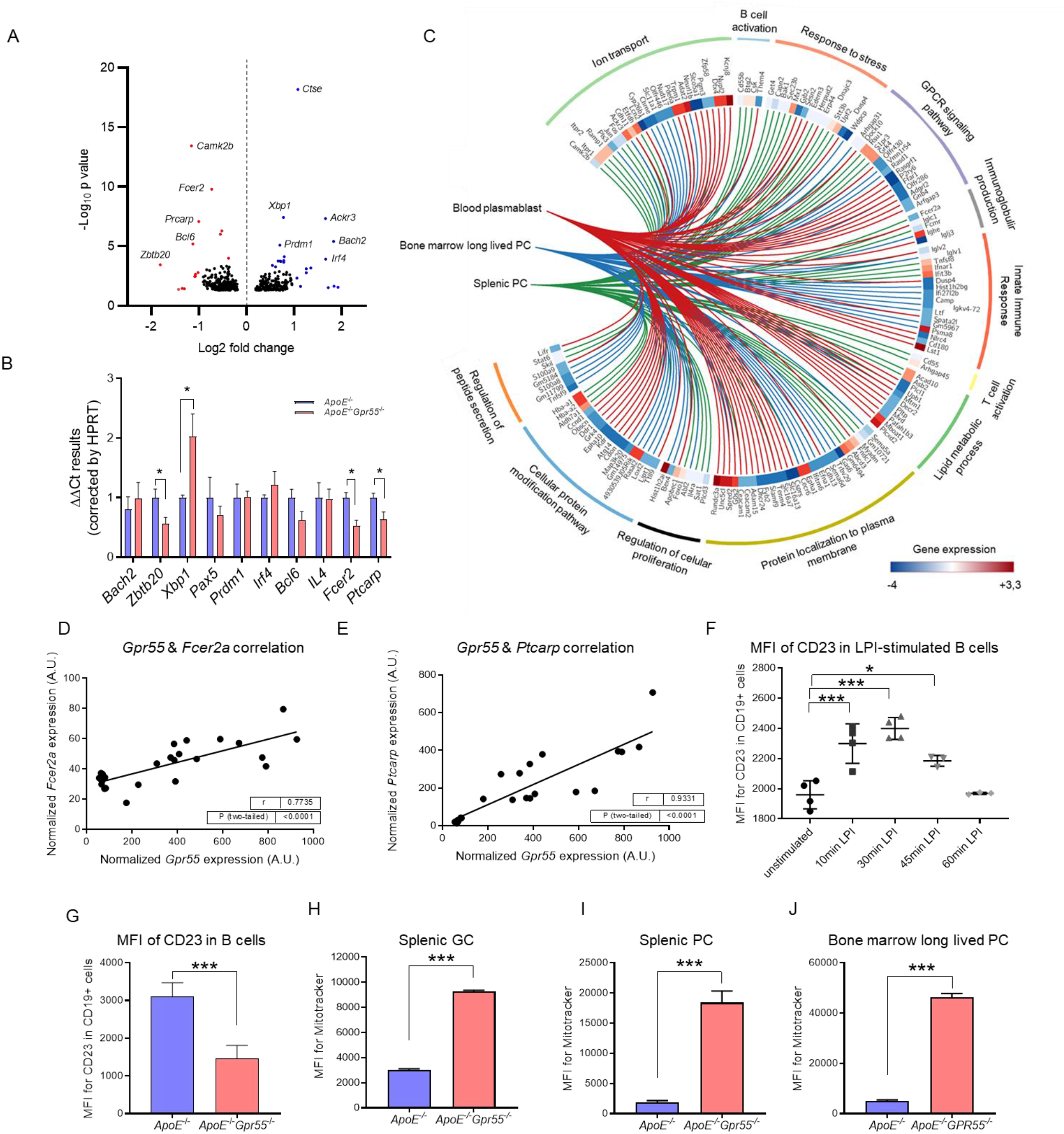
Impact of global *GPR55* deficiency on the B cell transcriptome, activation marker expression and mitochondrial content. (**A**) Volcano plot showing differentially expressed genes (DEGs) in sorted splenic CD19^+^ cells of *ApoE*^-/-^ and *ApoE^-/-^Gpr55^-/-^* mice after 4 weeks WD (adjusted p value 0.05, fold change ≥ log2 1.5-fold change). (**B**) Confirmation by qPCR of the main DEGs in splenic B cells of *ApoE*^-/-^ and *ApoE^-/-^Gpr55^-/-^* mice after 4 weeks WD. (**C**) Chord diagram showing the main regulated genes and pathways in splenic B cells, splenic PCs and circulatory plasmablasts, based on RNA-sequencing. (**D**) Splenic B cell *Gpr55* mRNA expression (qPCR) correlated with *Fcer2a* (CD23 encoding gene) mRNA expression. (**E**) Splenic B cell *Gpr55* mRNA expression (qPCR) correlated with *Ptcarpc* (LPAP encoding gene) mRNA expression. (**F**) Mean fluorescence intensity (MFI) of CD23 on splenic B cells measured by flow cytometry after LPI treatment (1 µM). (**G**) Flow cytometric analysis of CD23 MFI on splenic B cells of *ApoE^-/-^* and *ApoE^-/-^Gpr55^-/-^* mice after 4 weeks WD. (**H**) Mitochondrial content of splenic GC B cells. (**I**) Mitochondrial content of splenic PCs. (**J**) Mitochondrial content of bone marrow long lived PCs. Bulk RNA-sequencing analysis and qPCR validations (A-E) were performed with sorted cell from n=6 mice per group. Data shown in F-J were combined from 2 independent experiments (mean ± SEM; n=5-7 per group). Unpaired Student’s t-test was used to determine the significant differences *p < 0.05 and ***p < 0.001 vs the *ApoE^-/-^* group.

To further study the role of GPR55 in PC differentiation, we performed additional *in vitro* experiments. Splenic B cells isolated from *ApoE^-/-^* mice were stimulated with a cocktail of LPS, IFN-α, IL2, IL4 and IL5 for 72 h to trigger PC differentiation alone or in the presence of the GPR55-agonist LPI or the antagonist CID16020046, respectively. Co-incubation with LPI resulted in significantly more differentiated PCs compared to cells treated with LPS/IL2/IL5 alone, while adding the GPR55 antagonist prevented the LPS/IFN-α/IL2/IL4/IL5-induced induction of PC differentiation (Supplementary Figure 5A). This observation suggests that pro-inflammatory stimulation leads to endogenous GPR55 ligand production in splenic B cells, which triggers an autocrine receptor activation. In addition, GPR55 antagonism significantly augmented the mitochondrial content of *in vitro* differentiated PCs, which may reflect enhanced antibody secretion and cell stress (Supplementary Figure 5B). We subsequently performed confocal/STED imaging of unstimulated splenic PCs freshly sorted from *ApoE^-/-^* and *ApoE^-/-^GPR55^-/-^* mice, which uncovered a different morphology in *Gpr55*-deficient cells, with enlarged mitochondria and modified actin cytoskeleton (Supplementary Figure 5C and Supplementary videos).

Furthermore, we used flow cytometry to compare the mitochondrial content of splenic GC B cells, PCs and bone marrow long lived PCs between *ApoE^-/-^* and *ApoE^-/-^GPR55^-/-^* mice, which confirmed an increased mitochondrial content in all these subsets (Figure 4H-J). We may speculate that disturbed Ca^2+^ signaling due to the lack of GPR55 leads to ER stress and mitochondrial content.^31^ In summary, our findings support a crucial requirement for GPR55 signaling in regulating B cell activation and differentiation into PCs.

### B cell-specific *Gpr55* deficiency substantiates the role of GPR55 in modulating OSE antibody responses and atherosclerotic plaque development

To further address the specific contribution of B cell GPR55 signaling in atherosclerosis, we subsequently used a mixed bone marrow transplantation strategy, employing *µMT* as a model for mice lacking functional B cells, *Gpr55*-deficient bone marrow donors on wildtype (WT) background and *Ldlr*^-/-^ mice as recipients*. Gpr55^-/-^* bone marrow was mixed with *µMT* bone marrow in a 50/50 ratio before transplantation into lethally irradiated *Ldlr*^-/-^ recipients (Figure 5A). The B cell-specific deficiency was validated after 6 weeks recovery by qPCR of sorted circulating B cells, while T cells and myeloid cells exhibited normal expression levels of Gpr55 (Figure 5B). At the end point following recovery and 10 weeks WD feeding, the metabolic effects observed in mice with global GPR55 deficiency were lost, as we did not observe changes in body weight between groups and we could observe a slightly decreased plasma cholesterol in B-cell specific *Gpr55*-deficient mice compared to their respective controls (Figure 5C-D).

**Figure 5:**
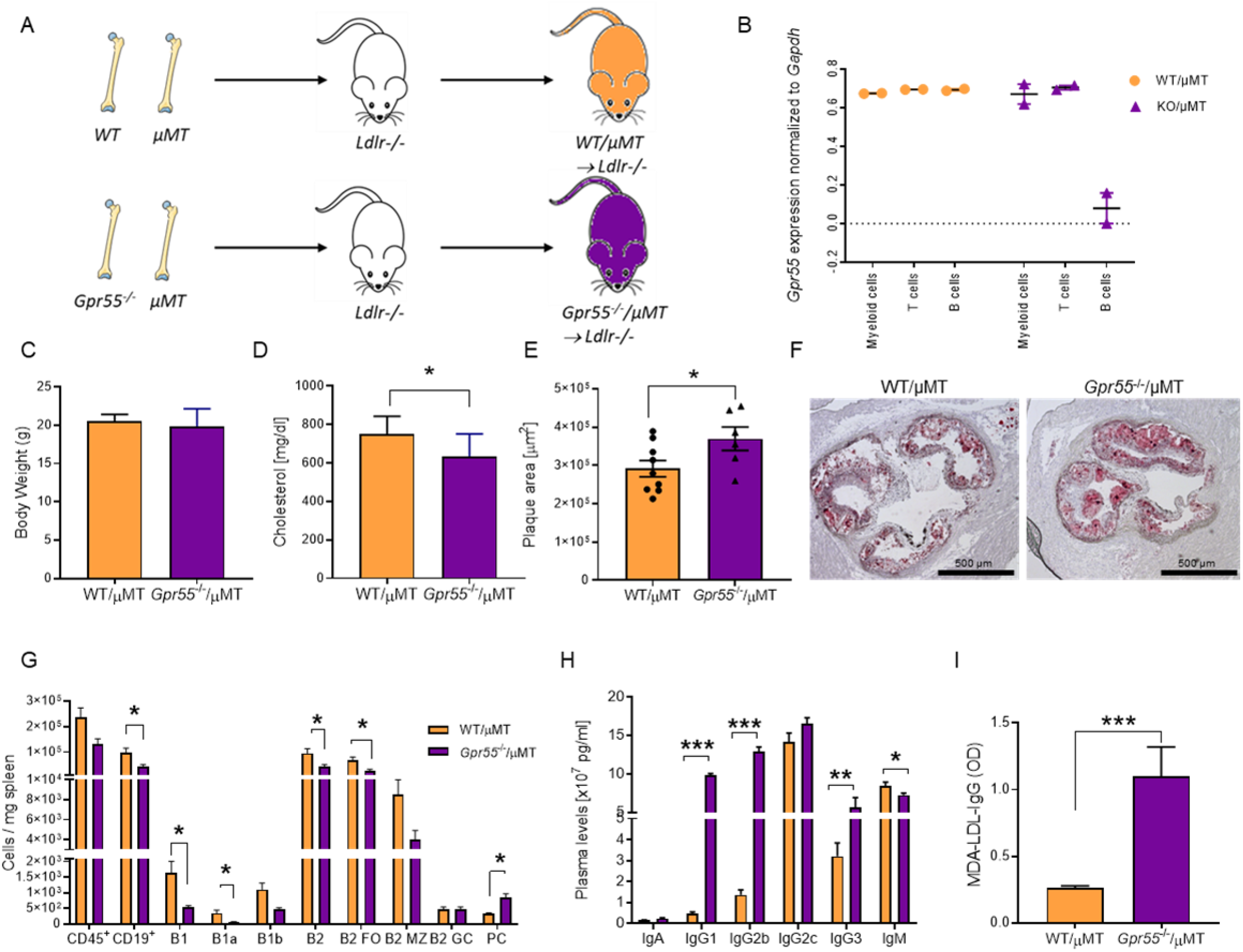
Impact of B cell-specific *Gpr55* deficiency on atherosclerotic plaque development and B cell responses during hypercholesterolemia. (**A**) Experimental design of the bone marrow transplantation to generate mixed bone marrow chimeras with B cell-specific *Gpr55* deficiency. Lethally irradiated *Ldlr^-/-^* mice received a 50/50 mixture of μMT marrow and *Gpr55^-/-^* or WT marrow cells. (**B**) Following 6 weeks recovery after the bone marrow transplantation, *Gpr55* expression was measured by qPCR in sorted blood myeloid cells (Cd11b^+^), T cells (CD3^+^) and B cells (B220^+^) to determine the B cell *Gpr55* depletion efficiency in *Gpr55^-/-^/µMT → Ldlr*^-/-^ mice. (**C**) Body weight and (**D**) total plasma cholesterol concentration after 10 weeks WD. (**E**) Plaque area in aortic roots of bone marrow chimeric mice after 10 weeks WD. (**F**) Representative Oil-Red-O stained aortic root sections used for plaque quantification. (**G**) Splenic B cell subsets assessed by flow cytometry. (**H**) Plasma Ig titers at the end of the experiment. (**I**) IgG antibodies against MDA-oxLDL in plasma after 10 weeks WD. Data were obtained in one bone marrow transplantation experiment (mean ± SEM; each point represents one animal; n=6-9). Unpaired Student’s t-test was used to determine the significant differences *p < 0.05 and ***p < 0.001 vs the indicated group

Mice with B cell specific GPR55 deficiency developed larger plaques compared to control mice transplanted with a mixture of WT/*µMT* bone marrow (Figure 5E-F), while B cell-specific *Gpr55* deficiency resulted in less splenic B1a cells and FO B2 cells, and PC counts were elevated (Figure 5G). The distinct profile in splenic B cell subsets of B cell-specific compared to global *Gpr55*-deficient mice indicates that the observed phenotype in global knockouts was not only due to B cell-dependent effects, but also modulated by T cell responses, such as Tfh interactions. Assessment of circulating antibody titers revealed slightly reduced plasma IgM titers, while IgGs were increased in B cells-specific *Gpr55*-deficient mice (Fig. 5H), which is in line with the proatherogenic phenotype. In particular, we observed significantly increased plasma titers of MDA-LDL-specific IgGs, which points to an involvement of specific responses to OSE under hypercholesterolemic conditions (Figure 5I).

### Adoptive transfer of WT B cells reverses the proatherogenic phenotype in global *ApoE^-/-^ GPR55^-/-^* mice

To substantiate the crucial importance of functional B cell GPR55 signaling in atherosclerosis, we performed an adoptive transfer experiment to rescue the proatherogenic phenotype in global *ApoE^-/-^Gpr55^-/-^* mice with WT B cells. We first treated *ApoE^-/-^* or *ApoE^-/-^Gpr55^-/-^* mice, respectively, with a B cell depleting antibody cocktail (Figure 6A-B) before transferring WT B cells. The adoptive transfer was sufficient to inhibit the proatherosclerotic phenotype, since *ApoE^-/-^Gpr55^-/-^* mice receiving WT B cells had comparable plaque sizes as observed in corresponding *ApoE^-/-^* control mice after 4 weeks WD (Figure 6C-D). However, the WT B cell transfer was not enough to entirely prevent the enhanced IgG response in *ApoE^-/-^Gpr55^-/-^* mice (Figure 6E), most likely because *Gpr55*-deficient plasmablasts and long-lived PC were not entirely depleted by the antibody cocktail, thereby still abundantly producing IgGs.

**Figure 6:**
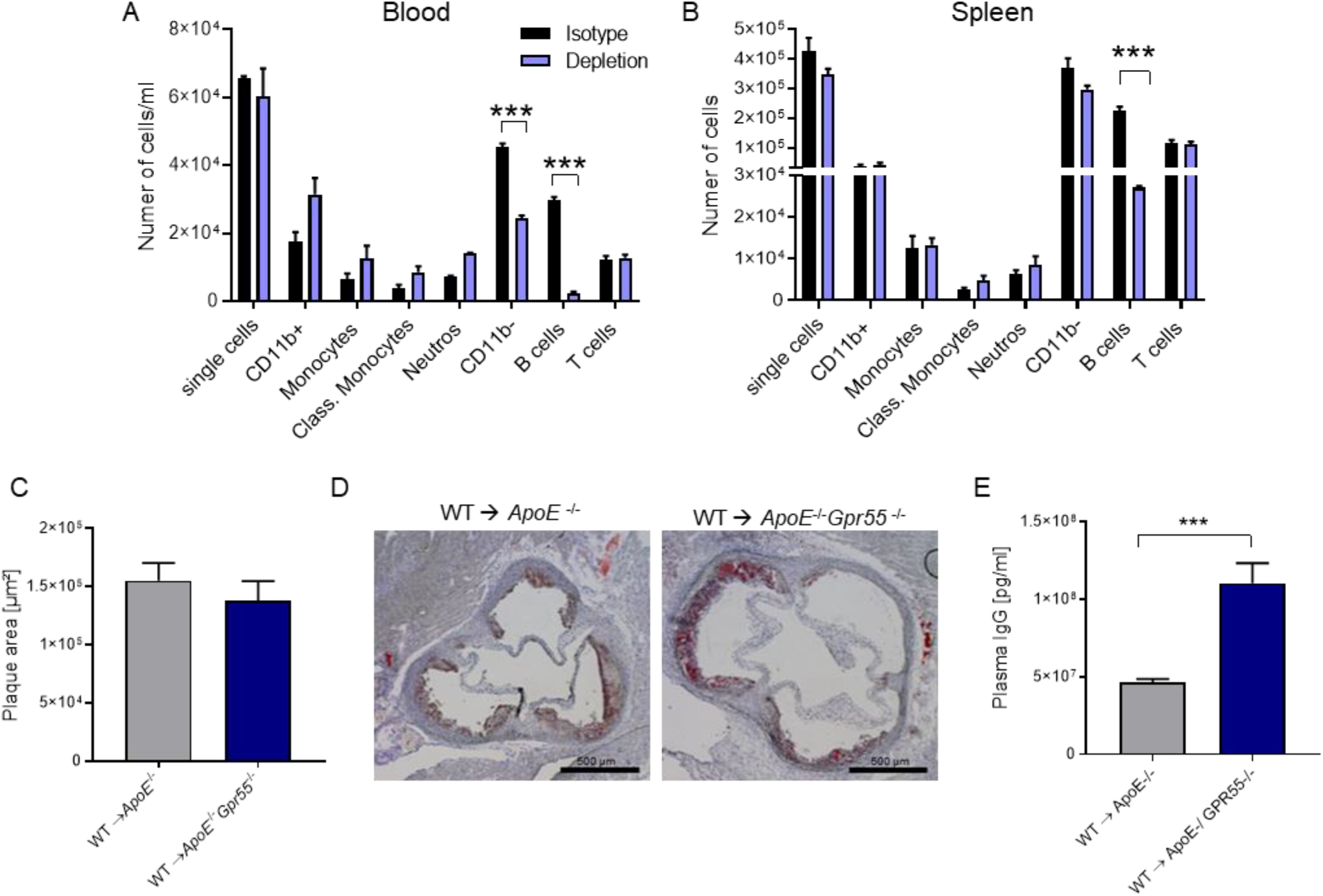
Impact of WT adoptive B cell transfer into global *Gpr55*-deficient mice on atherosclerotic plaque formation. (**A**) Circulating and (**B**) splenic leukocyte counts in *ApoE^-/-^Gpr55^-/-^* mice treated with the B cell depletion cocktail or control animals treated with the respective isotype antibodies (n=3 per group). (**C**) Aortic root plaque size and (**D**) representative oil red O-stained sections of B cell-depleted *ApoE^-^*^/-^ or *ApoE^-^*^/-^*Gpr55^-^*^/-^ mice receiving WT B cell adoptive transfer and subsequent WD feeding for 4 weeks. (**E**) Total plasma IgG levels after 4 weeks WD. Data were obtained in one experiment (mean ± SEM; n=5-7 per group). Unpaired Student’s t-test was used to determine the significant differences ***p < 0.001 vs indicated group.

## Discussion

In this study, we investigated the role of GPR55 signaling in atherosclerosis. Our findings show that GPR55 and its endogenous ligand LPI are regulated during atherogenesis. We report that GPR55 expression in mice is inversely correlated with plaque development and down-regulated in human unstable plaques compared to stable plaques, supporting the hypothesis that GPR55 signaling might play an atheroprotective role, both in a murine model and in human pathophysiology. We further provide direct evidence that global GPR55 expression plays a role in atherosclerosis and related metabolic dysregulation, as its general depletion resulted in more advanced plaque development, elevated plasma and liver cholesterol levels, together with increased body weight. Our liver metabolic data are somewhat controversial to recent findings in hepatocytes, where a pro-steatotic function of GPR55 signaling was documented by stimulation with the synthetic agonist O-1602, an effect that was reversed by the GPR55 antagonist CID16020046.^25^ Likewise, O-1602 increased serum triglyceride levels *in vivo*, which was prevented by co-administration of CID16020046.^25^ Moreover, circulating LPI levels and liver expression of GPR55 were up-regulated in patients with steatosis and nonalcoholic steatohepatitis (NASH). *In vivo* knockdown of *Gpr55* improved liver damage in mice fed either a high-fat diet or methionine-choline-deficient diet, and LPI also promoted the initiation of hepatic stellate cell activation by stimulating GPR55.^32^ A possible explanation might be that GPR55 signaling differentially affects lipid metabolism depending on the dietary condition, which deserves further investigation e.g. in mouse models with hepatocyte specific *Gpr55* deficiency.

The finding that GPR55 is particularly expressed by splenic B cells supports a central role for this receptor in regulating the homeostatic functions of this immune cell subset. It is well described that an imbalance of B cell homeostasis can cause devastating effects and tissue injury.^10, 13, 33, 34^ The hyperactivated and dysfunctional B cell state together with an exaggerated antibody production in hypercholesterolemic mice lacking functional GPR55 signaling fits well to the proatherogenic phenotype.^35–38^ B cell maturation and activation is precisely controlled by multiple factors, but begins with the binding of B-cell receptor (BCR) to a specific antigen,^39^ such as an OSE. The balance of activating and inhibitory signals in B cells is crucial for preventing an exaggerated adaptive immune response.^40^ Here, we present evidence that GPR55 signaling exerts a protective effect in B cells, while the lack of GPR55 is promoting an uncontrolled T and B cell interaction, leading to increased Tfh and circulating PC counts, together with elevated plasma BAFF levels, thereby cumulating in an excessive IgG production.

Having identified the importance of GPR55 in B cell activation, we further sought to identify the transcription factors, effectors or pathways affected by the lack of GPR55 signaling. Using bulk RNA-sequencing and GO enrichment analysis, we found that many genes and pathways involved in B and T cell activation, cellular response to stress and intracellular signal transduction were differentially regulated in *Gpr55* deficient B cells. By integrating all these pathways, we focused on the B cell activation- and maturation-related genes. Among the downregulated genes, we confirmed that *Fcer2a/CD23* and *Ptprc/LPAP* are positively correlated with *Gpr55* expression. The regulation of CD23 by GPR55 signaling was supported at the protein level by the reduced surface expression on CD19^+^ B cells of *ApoE^-/-^Gpr55^-/-^* mice and additional *in vitro* experiments with GPR55 agonists and antagonists. Even though CD23 has been extensively studied, its function is not fully understood. Previous studies using CD23 knockout mouse models suggested that CD23 is not critical for B and T cell maturation, but plays a regulatory function in the adaptive immune response.^41^ Liu *et al.* demonstrated that CD23 downregulates BCR signaling by inhibiting actin-mediated BCR clustering and B cell morphological changes.^27^ Moreover, a link between CD23 expression and IgG antibody response has been proposed.^42^ In particular, overexpression of CD23 resulted in significantly decreased IgG antibody responses. Hence, we may speculate that the decreased expression of CD23 in absence of GPR55 and thus lack of inhibitory signal for BCR signaling and IgG responses may contribute to the observed phenotype in our model. So far, not many external and internal signals and upstream signaling mechanisms that regulate the surface expression of CD23 have been identified. Our data suggest that GPR55 signaling represents such a B cell-intrinsic factor directly regulating CD23. The precise mechanism how GPR55 signaling is triggering CD23 expression remains to be addressed. Less is known regarding *Ptprc*/LPAP in B cells. *In vitro* stimulation of *Ptprc* mutant B cells led to normal differentiation to plasmablasts, but the cells failed to downregulate B220 expression, albeit it did not affect their functionality.^43^ Whether GPR55 signaling directly regulates LPAP and underlying mechanisms still needs to be clarified.

To better understand the mechanisms of dysregulated IgG production in mice lacking *Gpr55*, we focused on PCs, the main producers of antibodies. We present *in vitro* evidence that LPI directly triggers B cell maturation into PCs. Likewise, we found that the number of PCs was reduced when we antagonized GPR55 signaling. Moreover, PCs maturating *in vitro* under GPR55 blocking conditions had an increased mitochondrial content, which was supported by STED imaging of splenic PCs from *Gpr55*-deficient mice, revealing and altered mitochondrial and actin cytoskeleton structure. A link between mitochondrial dynamics, activation and differentiation of B cells has been previously described. Activated B cells exposed to high levels of BAFF increased glucose uptake and their mitochondrial mass, resulting in increased glycolysis and oxidative phosphorylation.^44, 45^ Although the mitochondrial dynamics in PCs have not been studied in detail, it is likely that IgG secretion in these cells is linked to increased metabolic rates and induces cellular stress responses.^44, 46–48^ The minor increase of IgM levels suggests that B cell GPR55 is not a key regulator for natural antibody production by B1 cells. This is contrasted by the role of another GPCR, namely CXCR4 on B1 cells, which controls IgM titers. Genetic deficiency of CXCR4 in B cells markedly exacerbated atherosclerosis, indicating the importance of B1 cell–derived IgM in atheroprotection.^49, 50^

The observed Ca^2+^ influx in response to LPI stimulation may play an important role in the regulation of cellular responses in PCs. Ca^2+^ signals affect a variety of intracellular processes that are central to cell-fate decisions in B cells, including mitochondrial physiology, apoptosis, secretion of transcription factors, cell migration, and Ig production.^51^ We may speculate that the lack of GPR55 signaling reduces intracellular/ER Ca^2+^ stores, which might lead to a defective BCR effector pathway activation, triggering a metabolic and biochemical state, that results in severe defects in B cell differentiation and function promoting atherogenesis. This, together with the downregulation of inhibitory signals such as CD23 may explain the unbalanced B cell immune response observed in mice lacking functional GPR55 signaling.

The bone marrow chimerism model to deplete GPR55 only in B cells confirmed the B cell-dependent atheroprotective effects of GPR55 signaling in atherosclerosis. However, some differences in the types of antibodies upregulated in mice with global or B cell-specific *Gpr55* deficiency exist, which may involve differences in T cell-dependent antibody responses in global *Gpr55* deficient mice. In fact, our gene expression analysis of sorted splenic immune cell subsets also indicates a strong GPR55 expression in T cells, which deserves further investigation in subsequent studies. Based on our mixed bone marrow chimera experiments, we can conclude that GPR55 signaling in B cells crucially regulates atherosclerosis. In support of a B cell-intrinsic role for GPR55 in atherogenesis, we could rescue the proatherogenic phenotype in global *Gpr55* deficient mice by WT B cell adoptive transfer to restore GPR55-expressing B cells. This strengthens our conclusion that intrinsic GPR55 signaling in B cells is necessary for their proper function and provides a negative regulatory signal to limit atheroprogression.

To conclude, our experimental *in vivo* and *in vitro* data, together with the confirmation of GPR55 expression and correlation with disease stage in human atherosclerosis, strongly support that targeting the GPR55-LPI axis should be investigated further as a possible strategy for preventing atheroprogression. Moreover, we uncovered a novel role for GPR55-LPI signaling in PC differentiation. The lack of this lipid signaling axis in B cells promotes an increased cellular stress with altered cytoskeleton remodeling, increased mitochondrial content and disturbed IgG production linked to the enhanced plaque formation. Hence, orphan receptors such as GPR55, which are activated by bioactive lipid mediators may represent an interesting novel therapeutic target to be further explored in the context of chronic inflammatory diseases such as atherosclerosis and beyond.

## Methods

### Global *Gpr55*-deficiency mouse model of atherosclerosis

*Gpr55*^-/-^ mice on C57BL/6J wildtype (WT) background were purchased from the European Mouse Mutant Archive (EM:02355), genotyped as previously reported^52^ and backcrossed with apolipoprotein E-deficient mice (*ApoE^−/−^*; strain # 002052, The Jackson Laboratory). Male and female *ApoE^-/-^* and *ApoE^-/-^Gpr55^-/-^*mice aged 7 to 9 weeks were fed with Western diet (WD; 0.15% cholesterol, Ssniff, TD88137) for either 4 or 16 weeks. At the end point, mice were anesthetized with ketamine/xylazine, and blood was obtained via cardiac puncture. Heart, aorta, spleen, and livers were harvested after PBS perfusion. All animal experiments were approved by the local Ethics committee (District Government of Upper Bavaria; License Number: 55.2-1-54-2532-111-13 and 55.2-2532.Vet_ 02-18-114) and conducted in accordance with the institutional and national guidelines. Sample size for the experiments was selected for achieve an *a priori* 85% statistical power for biologically significant difference (d=0.8).

### B cell-specific *Gpr55*-deficient bone marrow chimeras

Female *Ldlr^-/-^* mice, female *Gpr55^-/-^* mice on WT background and B cell-deficient (*μMT*) female mice (strain # 002288, The Jackson Laboratory) aged 6–8-weeks at the start of experimental regimes were used. *Ldlr^-/-^* mice were irradiated with 8 Gy gamma-radiation and reconstituted with 2 × 10^6^ mixed-bone marrow cells consisting of 50% *μMT* marrow and 50% *Gpr55^-/-^* or WT marrow, respectively. After irradiation mice were recovered for 6 weeks and then subjected to WD feeding for 10 weeks. After the 6-week-recovery time point, tail vein blood was collected to assess the transplantation efficiency. At the end point, organs were collected as described above.

### Adoptive transfer of B cells

B cells were isolated from spleens of WT female mice and enriched with a B cell isolation kit (Miltenyi Biotec MACS). The purity of isolated B cells was 96±0.8%. B cells were intraperitoneally (i.p.) injected into B cell-depleted female *ApoE^−/−^* or *ApoE^-/-^GPR55^-/-^* mice (1×10^6^ cells). Three days prior to the adoptive B cell transfer, recipient mice were treated with a cocktail of the following antibodies: B220 (clone RA3.3A1/6.1, ref: BE0067), CD19 (clone 1D3, Ref: BE0150) and CD22 (clone Cy34.1, BE0011), and 48h later anti-rat kappa (clone TIB216, Ref: BE0122, all antibodies from BioXCell; 150 µg i.p. per antibody). To test the B cell depletion efficiency, some mice were treated with a depletion antibody cocktail or isotype (IgG1, BE0083, BioXCell) and euthanized 72 h after the first injection to measure leukocyte counts in the blood and spleen by flow cytometry.

### Human material

Human carotid artery plaques were harvested during carotid artery endarterectomy (CEA) surgery, transported to the laboratory and snap frozen. Carotid tissue with an unstable/ruptured or stable plaque phenotype (as previously described^53, 54^ was cut in ∼50mg pieces on dry ice. The patients’ characteristics are summarized in Supplementary Table 4. The tissue homogenization was performed in 700 µl Qiazol lysis reagent and total RNA was isolated using the miRNeasy Mini Kit (Qiagen, Netherlands) according to manufacture’s instruction. RNA concentration and purity were assessed using NanoDrop. RIN number was assessed using the RNA Screen Tape (Agilent, USA) in the Agilent TapeStation 4200. Next, first strand cDNA synthesis was performed using the High-Capacity-RNA-to-cDNA Kit (Applied Biosystems, USA), following the manufacturer’s instructions. Quantitative real-time PCR was performed in 96 well plates with a QuantStudio3 Cycler (Applied Biosystems, USA), using TaqMan Gene Expression Assays (ThermoFisher, USA) for detection of the following expressed genes: *RPLPO* (Hs99999902_m1), *GPR55* Hs00271662_s1).

### Cholesterol and triglyceride measurement

Liver tissue (50-70 mg) was homogenized in 500 μl of 0.1 % NP-40 in PBS using a bead mill Tissue Lyser (Qiagen) and centrifuged for 2 min at 3000 x g to remove the insoluble material. Total plasma cholesterol and triglyceride concentrations were quantified in murine plasma and liver homogenates using colorimetric assays (CHOD-PAP and TAG-PAP Roche) according to the manufacturer’s protocol.

### Measurement of LPI by liquid chromatography-tandem mass spectrometry

Plasma samples were allowed to thaw on ice water, and 50 μl aliquots were transferred to 1.5 ml centrifugation tubes. After adding 300 μL of ice-cold ethyl acetate/hexane (9:1, v/v) containing the deuterated LPI as internal standards, tubes were vortexed for 30 seconds and immediately centrifuged for 15 minutes at 20 000 x g at 4° C. The upper organic phase was removed, evaporated to dryness under a gentle stream of nitrogen at 37° C, and reconstituted in 50 μl acetonitrile/water (1:1, v/v). Plasma concentrations of LPI were determined by liquid chromatography-multiple reaction monitoring as previously described and normalize to the total volume of supernatant.^55^

### Histological studies

Mouse hearts were isolated after perfusion with PBS and frozen in Tissue-tek (Sakura Finetek). Aortic roots were cut in 5-μm thick serial cryosections. The sections were stained with Oil-Red O for total plaque and lipid analysis and Masso’s trichrome for collagen in accordance with the guidelines for experimental atherosclerosis studies by the American Heart Association.^56^ Lesion size was analyzed in a blinded manner and quantified in 8 sections per heart, separated by 100 μm from each other, to cover the entire aortic root. Liver was embedded and frozen in Tissue-tek (Sakura Finetek). Pieces of liver tissue were cut in 5-μm thick serial cryosections. The sections were stained with Oil-Red O and we quantified the lipid deposition in liver using Image J.

For conventional immunofluorescence, aortic root sections were fixed for 5 min with 4% formalin and blocked for 1 h at room temperature (RT) with blocking buffer (PBS with 1% bovine serum albumin). Then the slides were incubated with Mac2 primary antibody overnight at 4° C for macrophage staining followed by anti-rat-AF488 for 2 h at RT and nuclear staining (Hoechst33342) for 5 min. Images were acquired using a Leica DM6000B fluorescence microscope equipped with a digital camera (DFC365, Leica). The percentage of macrophage plaque area was calculated with the Leica Application Suite LAS V4.3 software.

Splenic cryosections of 10-μm thickness were fixed for 5 min with 4% formalin and blocked for 1 h at RT with blocking buffer, then stained with directly conjugated antibodies overnight at 4° C. PNA-FITC (1:100) was used for identifying germinal center B cells, IgM-AF647 (1:250) for identifying MZ B cells and memory B cells, CD23-AF594 (1:100) for FO B cells, and Hoechst33342 for the nuclei staining. Digital images were acquired using a three-dimensional confocal laser scanning microscope (CLSM; Leica SP8 3X) equipped with a 100xNA1.40 oil immersion objective (Leica).

### *In situ* hybridization for *Gpr55*

To determine the presence of GPR55 in B cells we performed fluorescent *in situ* hybridization (FISH) by using a murine *Gpr55*-probe (VB6-3216575, Affymetrix). The splenic sections were cut and collected in RNAse free slides (slides cleaned with RNAse ZAP). RNase-free requirements were maintained up to post-hybridization steps. The sections were prefixed in 4% PFA for 5 min at RT and washed 3 times with PBS for 5 min. The tissue sections were treated with pre-warmed 10 μg/ml proteinase K (diluted in PBS) for 5 min at RT, followed by post-fixing with 100% ethanol for 1 min. Then, the sections were washed 3 times for 10 min with PBS and transferred to an RNAse free chamber. The ViewRNA™ Cell Plus Assay-Kit (Invitrogen) was used for target probe hybridization by incubating the sections at 40° C for 2 h. The hybridization probe was mixed with probe set diluent to a final concentration of 5 μg/ml. Following the hybridization step, the sections were washed three times with the wash buffer at RT for 5 min. Signal amplification was performed by incubating with the preamplifier mix for 30 min at 40° C, washing and incubating with the amplifier mix for 1 h at 40° C. After washing, the sections were incubated for 1 h at 40° C with the working label probe mix. Following the in-situ hybridization for the *Gpr55*-probe, the sections were washed, incubated with blocking buffer at RT for 1 h and stained with PNA-FITC, IgM-AF647 and CD23-AF594 antibodies as described above.

### Flow cytometry

Splenic single cell suspensions were obtained by mashing the spleens through a 70-µm cell strainer. Femurs were centrifuged at 10,000 x g for 1 min after exposure of the distal metaphysis to collect the bone marrow cells. Splenic, bone marrow and blood erythrocytes were lysed with ammonium-chloride-potassium (ACK, NH_4_Cl (8,024 mg/l), KHCO_3_ (1,001 mg/l) EDTA.Na_2_·2H_2_O (3.722 mg/l) buffer for 10 min at RT. For staining of leukocytes infiltrated into the aorta, the vessels were excised from the aortic arch to iliac bifurcation, and digested with collagenase IV and DNase I at 37° C at 750 rpm for 40 min as previously described^57^ and filtered through a 30-µm cell strainer. Blood, splenic, bone marrow, and aortic single cells were blocked for 5 min with Fc-CD16/CD32 antibody and then stained for 30 min in the dark at 4° C with antibodies (Supplementary Table 1) to identify myeloid and lymphoid cell subsets. After gating for living singlets and CD45^+^, the CD11b^+^ myeloid subsets were further gated as following: CD115^+^Ly6G^-^ (monocytes), F4/80^+^ (macrophages), CD115^-^Ly6G^+^ (neutrophils). The CD11b^-^ lymphoid population was divided into CD3^+^ (T cells) and CD19^+^ (B cells). B cell subsets were further identified as B220^low^CD23^-^CD5^+^ (B1a), B220^low^CD23^-^CD5^-^ (B1b), B220^high^CD23^-^IgM^+^ (MZ), B220^high^CD23^-^IgM^int^IgD^+^ (FO), B220^high^CD23^+^IgM^int^PNA^+^GL7^+^ (GC). PCs were gated as CD19^-^CD138^+^CD23^-^IgM^-^IgD^-^ and splenic Tfh were gated as CD3^+^CD19^-^CXCR5^+^PD1^+^. The circulating plasmablasts were identified as CD19^+^CD38^+^CD138^+^ and long-lived bone marrow PCs as CD19^-^CD138^+^. For mitochondrial staining in splenic and bone marrow B cells, the cells were incubated with prewarmed (37° C) staining solution containing MitoTracker (ThermoFisher) at a concentration of 50 nM for 30 min. After removing the mitochondrial staining solution by centrifugation, the cells were stained with specific antibodies to identify the different B cell subpopulations. Flow cytometry data were acquired on a BD FACSCanto II flow cytometer (BD Biosciences) or Fortessa LSR (BD Biosciences) and analyzed with FlowJo v10.2 software (Tree Star, Inc).

### Flow cytometric sorting of splenic B cells

Single cell suspensions obtained from *ApoE^-/-^* and *ApoE^-/-^Gpr55^-/-^* spleens were sorted (FACs ARIA) as CD19^+^ total B cells or B220^high^CD23^+^IgM^int^PNA^+^GL7^+^ GC cells. Circulating plasmablasts were gated asCD19^+^CD38^+^CD138^+^, and bone marrow long-lived PCs asCD19^-^CD138^+^. The sorted cells were deep frozen in 2x TCL buffer plus (Qiagen) plus 1% beta-mercaptoethanol.

### RNA-sequencing and analysis

Splenic PC, bone marrow long lived PCs, and circulating plasmablasts from 6 donor mice per group (*ApoE^-/-^* versus *ApoE^-/-^Gpr55^-/-^* mice) were sorted using a BD FACSAria III Cell Sorter (BD Biosciences) as described above and bulk RNA-sequencing was performed using the prime-seq protocol.^26^ A step-by-step protocol can be found on protocols.io (dx.doi.org/10.17504/protocols.io.s9veh66). Briefly, 10,000 sorted cells were lysed in 100 µL of RLT+, 1% beta-mercaptoethanol and 50 µL of lysate was used for RNA-sequencing. In case that less than 10,000 cells could not be obtained, all sorted cells were used. The samples were proteinase K and DNase I digested and then cDNA synthesis was performed using uniquely barcoded oligodT primers. Samples destined for the same library were pooled and pre-amplification was then performed using 11-14 cycles, depending on the initial input per library. The cDNA was quantified using the PicoGreen dsDNA assay kit (Thermo Fisher, P11496) and qualified using the Bioanalyzer HS DNA chip (Agilent, 5067-4626). Libraries were then constructed with the NEB Next Ultra II FS kit (E6177S, NEB) using the prime-seq specifications. The libraries were quantified and qualified using the HS DNA chip on the Bioanalyzer and sequenced on an Illumina Hiseq 1500 at an average depth of 12.7 million reads per sample. The reads were demultiplexed using deML and then filtered, mapped to the mouse genome (mm10, GRCm38), and counted using zUMIs (version 2.5.5) with STAR.^58^ Differential gene expression analysis was performed using DESeq2 and then the characterization of the molecular functions or pathways in which differentially expressed genes (DEGs) are involved was performed using gene set enrichment analysis (GSEA). Resources used were Gene Ontology, Bioconductor, Gorilla and DAVID.

### Quantitative real-time polymerase chain reaction

Aortas were homogenized and lysed for extraction of total RNA (peqGold Trifast and Total RNA kit, Peqlab). RNA from sorted splenic B cells was also isolated using the same kit. After reverse transcription (PrimeScript RT reagent kit, Clontech), real-time polymerase chain reaction was performed with the 7900HT Sequence Detection System (Applied Biosystems) using the KAPA PROBE FAST Universal qPCR kit (Peqlab). Primers and probes were purchased from Life Technologies. Target gene expression was normalized to *Hprt* (hypoxanthine-guanine phosphoribosyltransferase) and presented as relative transcript level (2−^ΔΔCt^). For comparison of *Gpr55* expression, *Gapdh* (glyceraldehyde-3-phosphate dehydrogenase) was used as additional housekeeping control for normalization (Supplementary Table 2).

### Digital droplet PCR

For performing the digital droplet-PCR (ddPCR) we prepared the reaction mixture by combining a 2× ddPCR Mastermix (Bio-Rad), 20× primer, and probes (final concentrations of 900 and 250 nM, respectively; Integrated Data Technologies) and template in a final volume of 20 μl (Supplementary Table 3). Then, each ddPCR reaction mixture was loaded into the sample well of an eight-channel disposable droplet generator cartridge (Bio-Rad). A volume of 60 μl of droplet generation oil (Bio-Rad) was loaded into the oil well for each channel. The cartridge was placed into the droplet generator (Bio-Rad), and the droplets were collected and transferred to a 96-well PCR plate. The plate was heat-sealed with a foil seal, placed on the thermal cycler and amplified to the end-point (40 cycles). After PCR, the 96-well PCR plate was loaded on the droplet reader (Bio-Rad). Analysis of the ddPCR data was performed with QuantaSoft analysis software (Bio-Rad).

### Immunoglobulin measurement in plasma

To measure Ig isotypes and subclasses in plasma, the Antibody Isotyping 7-Plex Mouse ProcartaPlex™ Panel was performed with a MAGPIX luminex reader (ThermoFisher). For total IgG and IgM quantification, single ELISA kits were used (ThermoFisher).

Anti-MDA specific IgGs were measured by coating MaxiSorp plates (NuncTM, city, Roskilde, Denmark) with purified, mouse-derived delipidated apolipoprotein MDA for 1 h at 37° C. After being washed, all wells were blocked for 1 h with 2% bovine serum albumin (BSA) in a phosphate-buffered solution (PBS) at 37° C. Mouse samples were also added to a non-coated well in order to assess individual non-specific binding. After six washing cycles, 50 µl/well of the alkaline phosphatase-conjugated anti-mouse IgG was added (Sigma-Aldrich, St Louis, MO), it was diluted at 1:1000 in a PBS/BSA 2% solution, and this was added and incubated for 1 h at 37° C. After washing six more times, phosphatase substrate p-nitrophanylphosphate disodium (Sigma-Aldrich) dissolved in a diethanolamine buffer (pH 9.8) was added and incubated for 30 min at 37° C. Optical density (OD) was determined at 405 nm (Filtermax 3, Molecular DevicesTM, San Jose, CA) and each sample was tested in duplicate. Corresponding non-specific binding was subtracted from mean OD for each sample.

### *In vitro* PC differentiation

Splenic B cells were isolated from *ApoE*^-/-^ mice by meshing the spleen and subsequent entrichment for CD19^+^ cells using a Miltenyi kit. For PC *in vitro* differentiation, the isolated B cells were seeded in 24-well plates and treated with a cocktail of LPS (0,5 ng/ml), IFN-α (2 ng/ml), IL2 (5 ng/ml), IL-4 (5 ng/ml), and IL5 (2 ng/ml) for 7 days, adding fresh medium with the stimulation cocktail every second day, based on an optimized protocol according to previous publications.^59, 60^ In some conditions, the stimulation cocktail was combined with LPI or CID16020046. After 7 days, cells were collected and used for flow cytometric analysis of the total number of PCs, gated as viable single CD45^+^CD11b^-^CD19^+^CD138^+^ cells.

### Confocal/STED imaging of splenic PCs

Splenic B cells were isolated from *ApoE*^-/-^ and *ApoE*^-/-^*Gpr55*^-/-^ mice, seeded into glass-bottom chamber 8-well slides (Ibidi) and differentiated into PCs as described above. The differentiated cells were incubated with mitotracker (Invitrogen) for 30 min at 37° C and afterwards washed, fixed in 2% paraformaldehyde (PFA) for 10 min and permeabilized with 0.1% Triton X-100 in PBS/BSA 1% for 30 min at RT. The slides were subsequently stained with F-actin for 30 min at RT, followed by incubation for 5 min with Hoechst33342 (ThermoFisher Scientific) and embedded with ProLong Diamond Antifade Mountant (ThermoFisher). Digital images were acquired using a three-dimensional CLSM combined with STED (Leica SP8 3X) equipped with a 100xNA1.40 oil immersion objective (Leica). Optical zoom was used where applicable. For fluorescence excitation, a UV laser (405 nm) was used for excitation of Hoechst and a tunable white light laser for selective excitation of other fluorochromes (Alexa488, Alexa594 and Alexa647). Depletion was performed at 592 nm and 775 nm for AlexaFluor488 and AlexaFluor594, respectively. Images were collected in a sequential scanning mode using hybrid diode detectors to maximize the signal collection and reduce the background noise and overlap between the channels. All data were acquired in three dimensions and voxel size was determined according to Nyquist sampling criterion. Image reconstructions were performed using the LAS X software package v.3.0.2 (Leica) and deconvolution was applied in combination with the Huygens Professional software package v.19.10 (Scientific Volume) using the unsupervised CMLE (for CLSM) algorithm. Visualization of 3D images is provided as a video showing the merged fluorescent channels from different visual perspectives (Supplemental videos). To this end, 3D rendering was performed using the LAS X 3D software and visual perspective was automatically selected by the software to provide the best visualization.

### Ca^2+^ assay

Ca^2+^ responses and CD23 surface expression was measured in murine blood after cell stimulation with LPI for 30 s. Briefly, the cells were stained with an antibody cocktail to identify cell subpopulations, washed and then incubated with FLIPR, Calcium 5 Assay Kit (Molecular devices) for 1 h at 37° C. The samples were subsequently measured by flow cytometry to determine the mean fluorescence intensity (MFI) of intracellular Ca^2+^ or surface CD23 expression, respectively.

## Statistical Analysis

Statistical analyses were performed using GraphPad Prism 7 software. To test for Gaussian distribution, D’Agostino Pearson omnibus or Shapiro–Wilk normality test was applied. Outliers were determined by Grubbs’ test (alpha 0.05). After testing homogeneity of variances via *F* test, Student’s *t* test was used for normally distributed data with equal variances. For heteroscedastic data, Welch correction was applied. Mann–Whitney *U* test was performed if normality test failed. Bivariate correlations involving *Gpr55* expression were analyzed by Spearman’s rank correlation test, to avoid assumptions on data distribution. All data are shown as mean ± SEM. A 2-tailed p<0.05 was considered statistically significant.

## Acknowledgements

We are deeply grateful to Maria Aslani for her advice in transcriptomic data visual presentation and thank Diana Wagner, Yvonne Jansen, Rodrigo Carrasco-Leon, Blanca Dufner and Silviya Wolkerstorfer for their excellent technical support.

## Author contributions

R. Guillamat-Prats, D. Hering and M. Rami designed and performed experiments and analyzed and interpreted data under S. Steffens’ supervision. C. Härdtner and I. Hilgendorf designed the B-cell bone marrow transplant experiment, and I. Hilgendorf gave critical input for the entire study design. D. Santovito provided expertise for confocal/STED imaging and statistical analysis. P. Rinne contributed to the histological analysis. L. Bindila performed lipidomic measurements. S. Pagano and N. Vuilleumier determined MDA-oxLDL antibodies in plasma. M. Hristov conducted cell sorting. A. Janjic and W. Enard performed the bulk RNA-sequencing and the clean-up of the obtained data. S. Schmid and L. Maegdefessel determined *GPR55* expression in human plaques.

A. Faussner guided the Ca^2+^ assays and provided critical input to the study. C. Weber provided scientific infrastructure and advice for the study design. R. Guillamat-Prats and S. Steffens designed the study and wrote the manuscript. All authors contributed to the final manuscript editing and approved the submitted version.

## Supplementary figures

**Supplementary Figure 1:**
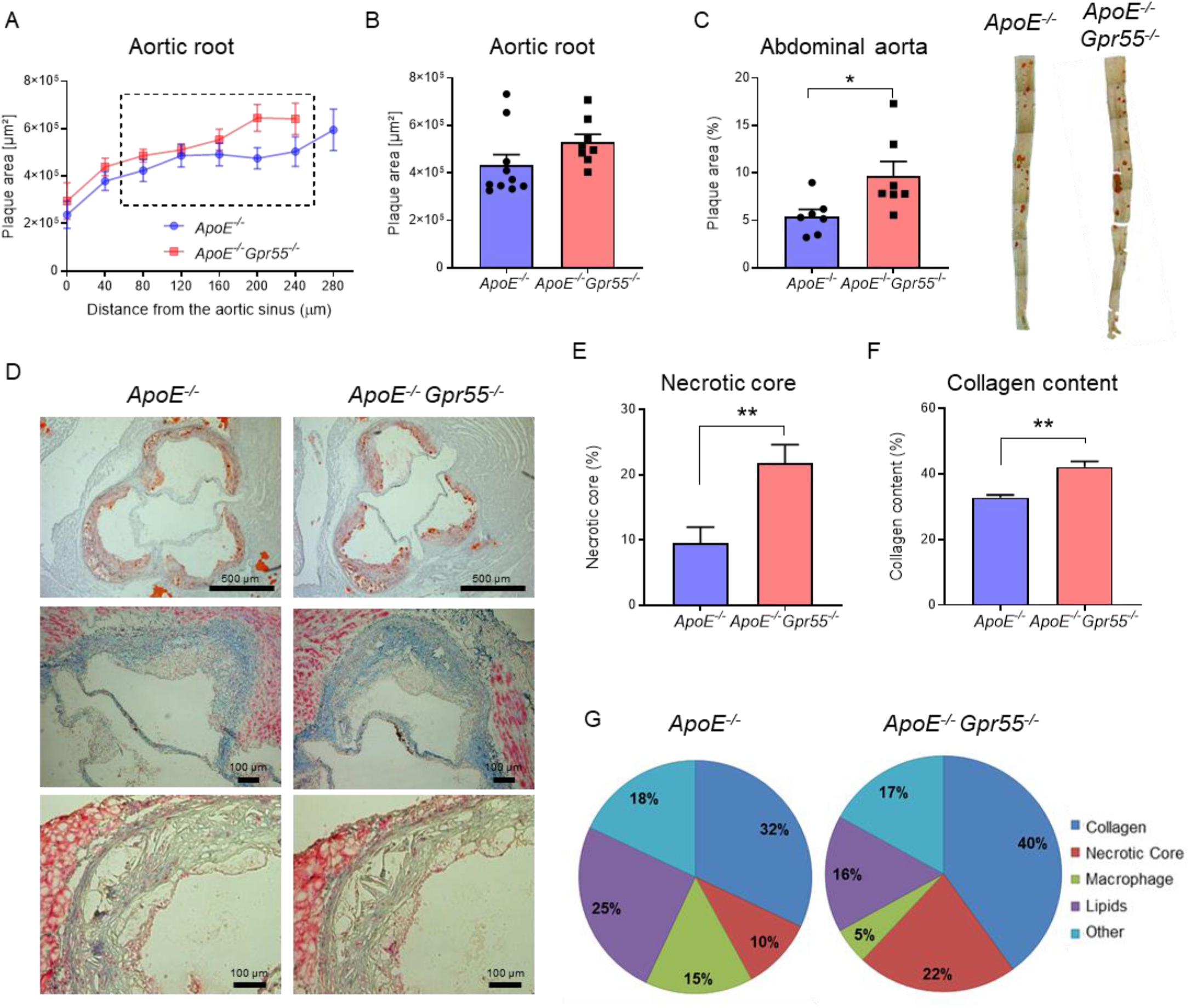
Impact of global GPR55 deficiency on advanced stage of atherosclerosis. (**A**) Plaque area per section and (**B**) average plaque area (calculated from sections 80-240 µm distance of the aortic root origin) after 16 weeks WD. (**C**) Plaque area within descending thoraco-abdominal aortas after 16 weeks WD, determined by *en face* preparations stained with Oil-red-O, and calculated as percentage of plaque per total vessel area. (**D**) Representative aortic root plaque images stained with Oil-red-O (upper) and Masson staining’s (middle and lower images) for lipid or collagen content, respectively, and quantification of the necrotic core area after 16 weeks WD. (**E**) Necrotic core and (**F**) collagen content per total plaque area were calculated on 4 middle aortic root sections per mouse heart. (**G**) Relative plaque composition of main components in advanced plaques after 16 weeks WD. Data were combined from 3 independent experiments (**A**-**C**, n=12-15 female mice per group and **E**-**G**, n=6-8 female mice per group; mean ± SEM). Unpaired Student’s t-test was used to determine the significant differences **p < 0.01 vs the indicated group.

**Supplementary Figure 2:**
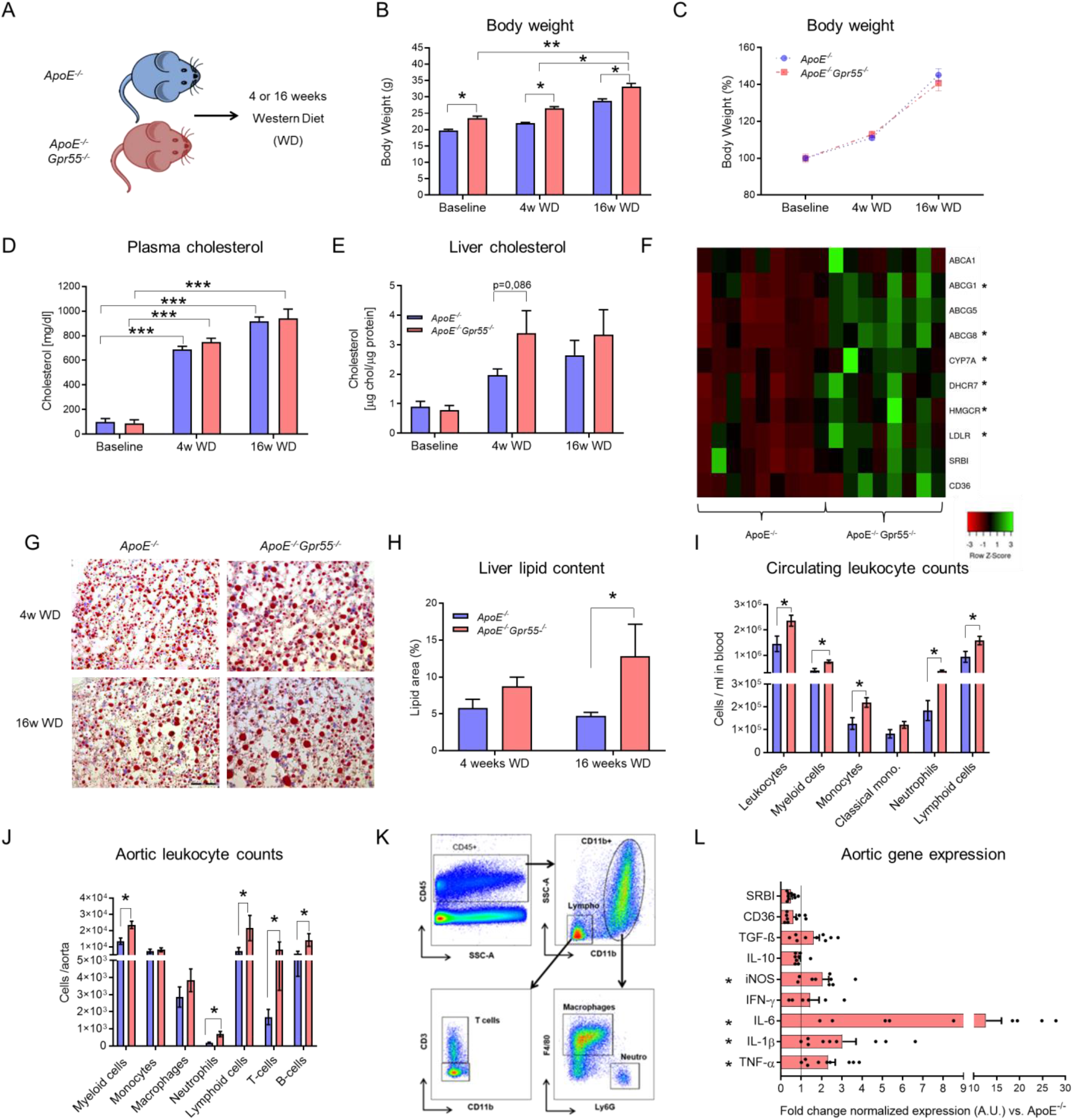
Global *Gpr55* deficiency leads to increased metabolic dysfunction and inflammation during hypercholesterolemia. (**A**) Experimental scheme of the WD treatment and color code for *ApoE^-/-^* and *ApoE^-/-^ Gpr55^-/-^* groups. (**B**) Average body weight of *ApoE^-/-^* vs *ApoE^-/-^ Gpr55^-/-^* female mice (n=12 for baseline, n=14-18 for 4 and 16 weeks WD) from different experimental end points (baseline, 4 or 16 weeks WD). (**C**) The body weight gain per mouse was followed for the entire duration of a 16 weeks WD experiment, and calculated as percentage of the starting body weight. (**D**) Total plasma cholesterol concentrations at baseline, 4 and 16 weeks WD from *ApoE^-/-^* and *ApoE^-/-^ Gpr55^-/-^* female mice. (**E**) Liver cholesterol concentrations, normalized to the total protein content per lysate, in *ApoE-/-* and *ApoE^-/-^ Gpr55^-/-^* mice at baseline, 4 and 16 weeks WD (D-E, n= 6 for baseline, n=10-12 for 4 and 16 weeks WD). (**F**) Heatmap showing the relative liver mRNA expression of several genes involved in cholesterol metabolism for *ApoE^-/-^* vs *ApoE^-/-^ Gpr55^-/-^* after 4 weeks WD. Each column represents one mouse (n= 8-9 per group). (**G**) Histological liver sections stained with oil-Red-O for lipid droplet quantification (**H**), calculated per tissue area, after 4 and 16 weeks WD in *ApoE^-/-^* vs *ApoE^-/-^ Gpr55^-/-^* mice (n=5-6 mice per group). (**I**) Circulating leukocyte subsets per ml of blood in *ApoE^-/-^* vs *ApoE^-/-^ Gpr55^-/-^* after 4 weeks WD were determined by flow cytometry. (**J**) Aortic leukocyte subsets in *ApoE^-/-^* vs *ApoE^-/-^ Gpr55^-/-^* mice after 4 weeks WD (I-J, n=10-12). (**K**) Flow Cytometry gating strategy used for identification of aortic leukocyte subsets (**L**) Aortic mRNA expression of representative key pro- and anti-inflammatory genes in *ApoE^-/-^* vs *ApoE^-/-^ Gpr55^-/-^* mice after 4 weeks WD (n=9-10). Data were combined from 2 independent experiments (mean ± SEM; data from female mice are shown). One way-ANOVA followed by post-hoc Newman–Keuls multiple-comparison test was used to evaluate the significant differences *p<0.05, ** p<0.01, ***p<0.001 vs the indicated group.

**Supplementary Figure 3.**
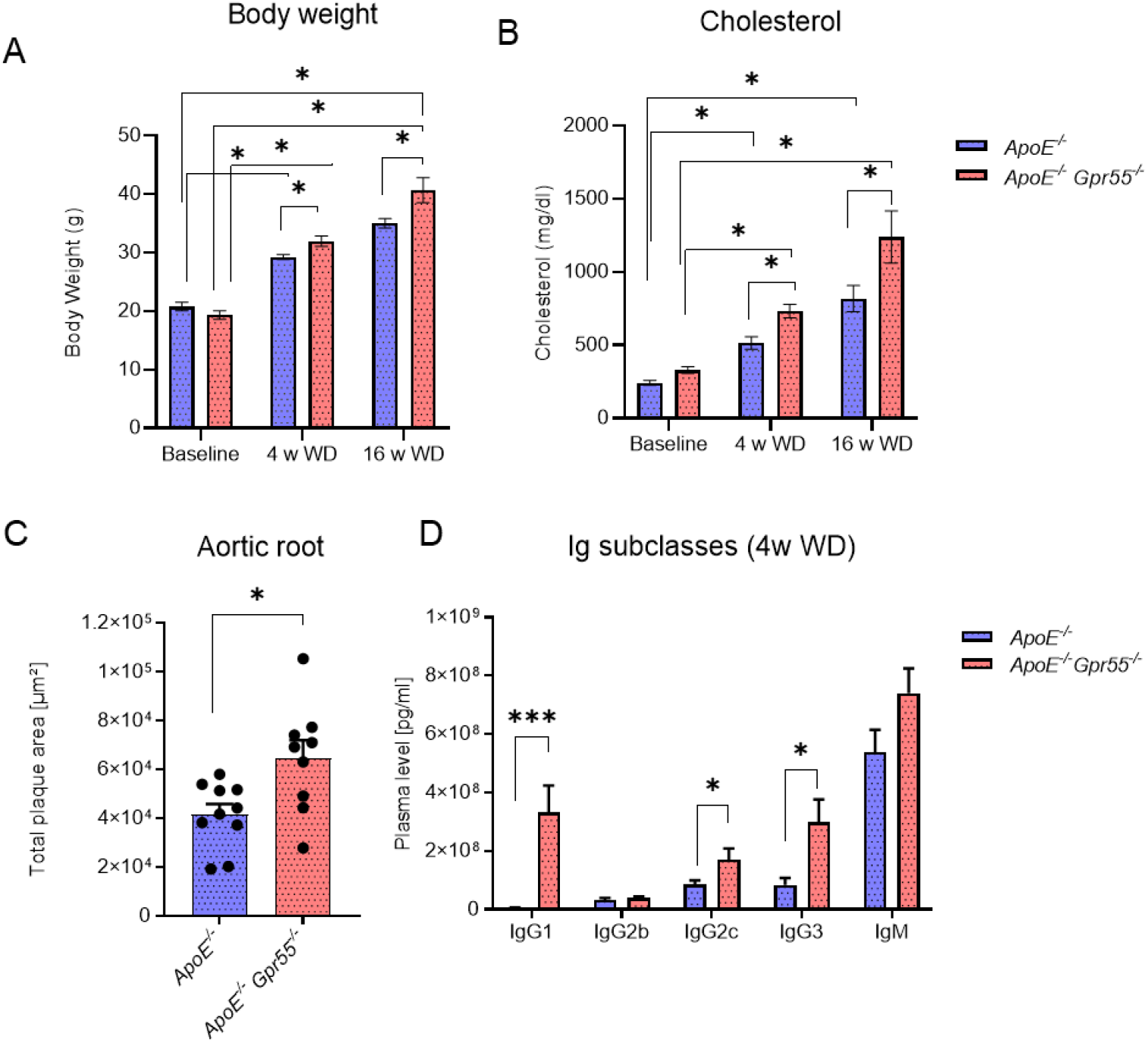
: Effects of global *Gpr55* deficiency in male mice (**A**) Average body weight of *ApoE^-/-^* vs *ApoE^-/-^ Gpr55^-/-^* mice from different experimental end points (baseline, 4 or 16 weeks WD). (**B**) Total plasma cholesterol concentrations at different time points (Baseline, 4 weeks WD, and 16 weeks WD) from *ApoE^-/-^* and *ApoE^-/-^ Gpr55^-/-^* mice. (**C**) Average plaque area in Oil-red-O-stained aortic root sections (calculated from sections 80-240 µm) after 4 weeks WD in mice. (**D**) Plasma Ig titers after 4 weeks WD. Data are mean ± SEM; A-D, n= 10-12 male mice per group. For panels A and B, one way-ANOVA followed by post-hoc Newman–Keuls multiple-comparison test was used to evaluate the significant differences *p<0.05, ***p<0.001 vs the indicated group. In panel C and D, unpaired Student’s t-test was used to determine the significant differences **p < 0.01 vs *ApoE^-/-^*.

**Supplementary Figure 4:**
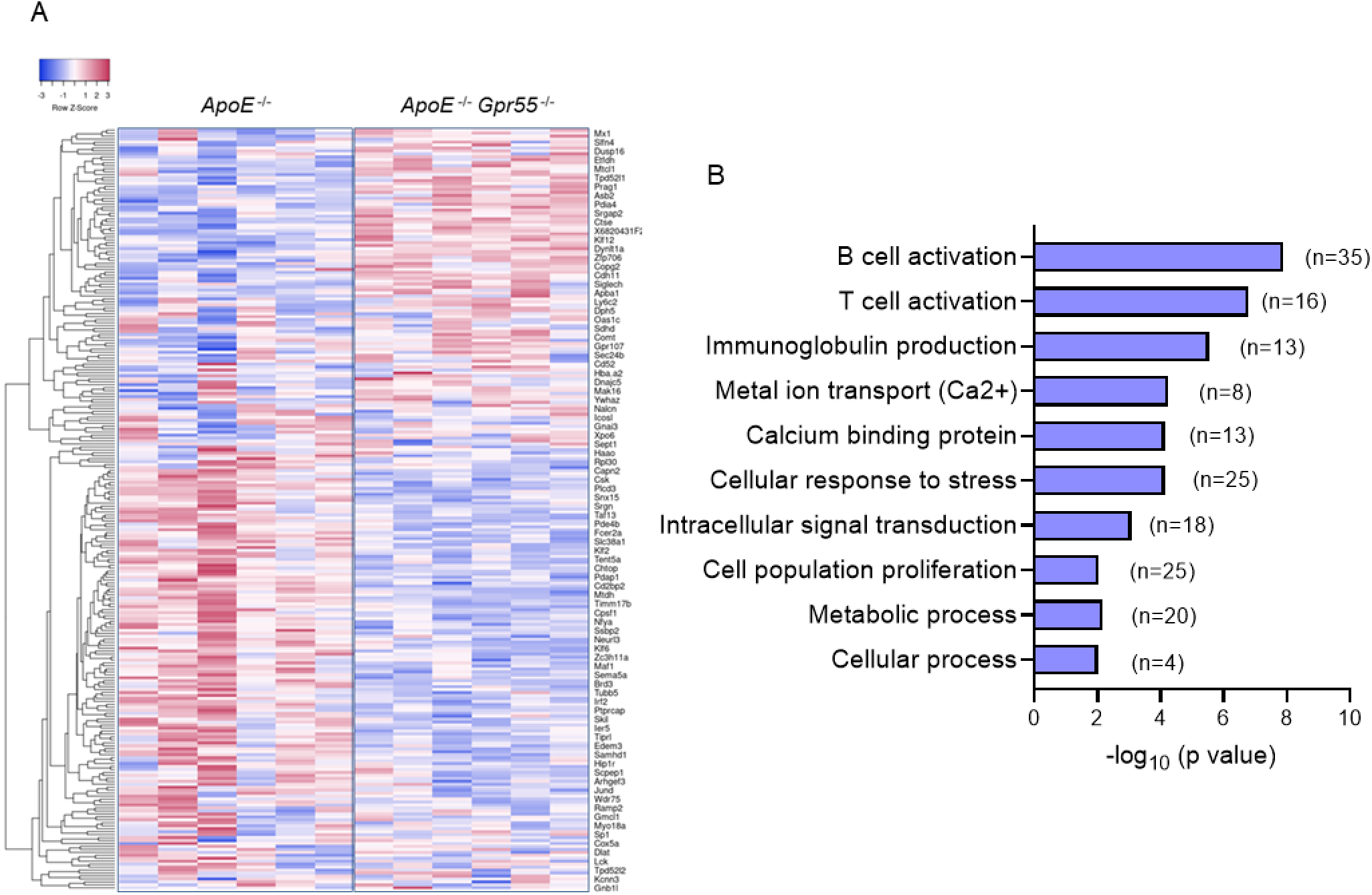
RNA-sequencing of total splenic B cells from *ApoE^-/-^* vs *ApoE^-/-^ Gpr55^-/-^* mice. (A) Heatmap of the differentially expressed genes between splenic B cells (CD19^+^) isolated from *ApoE^-/-^* and *ApoE^-/-^Gpr55^-/-^* mice after for 4 weeks WD (adjusted p value 0.05, fold change ≥ log2 1.5 fold change). (B) Main regulated pathways based on gene ontology (GO) enrichment analysis, bars represent the p value significance; numbers in brackets indicate the number of genes regulated in each pathway.

**Supplementary Figure 5.**
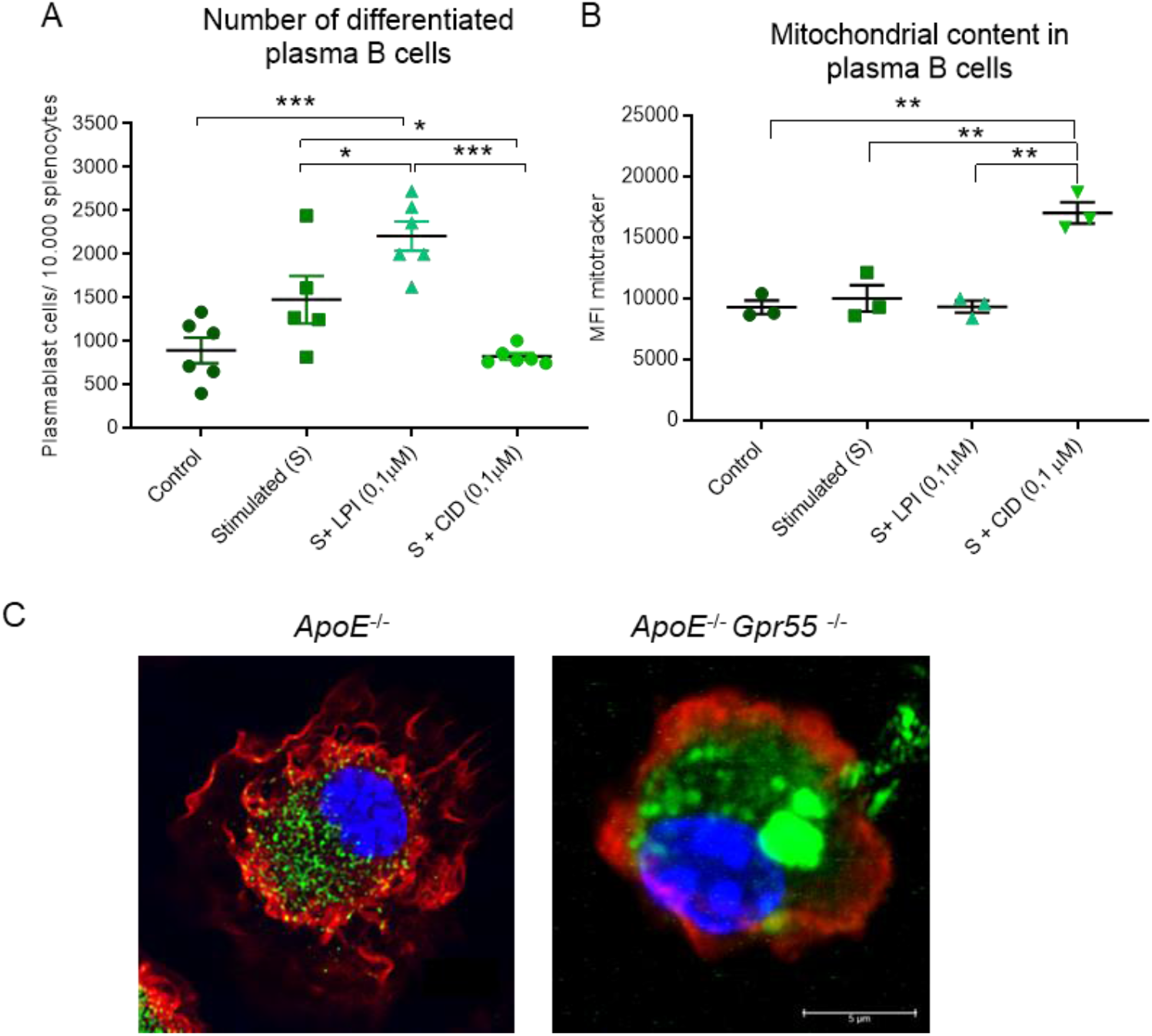
: Plasma B cell differentiation and characterization (**A, B**) *In vitro* differentiation of splenic B cells into PCs alone or in presence of GPR55 ligand LPI or GPR55 antagonist CID16020046. After 7 days, PCs were quantified by flow cytometry gating for CD45^+^CD11b^-^CD19^+^CD138^+^ cells. (**B**) Mitochondrial content of differentiated PCs was determined by flow cytometry using Mitotracker, data are expressed as MFI (each dot represents a separate *ApoE^-/-^* mouse used for B cell in vitro differentiation). (**C**) Representative confocal/STED images of differentiated PC from an *ApoE^-/-^* or *ApoE^-/-^Gpr55^-/-^* stained with Mitotracker (mitochondria, green), F-actin (red) and nuclei (blue). One way-ANOVA followed by post-hoc Newman–Keuls multiple-comparison test was used to evaluate the significant differences *p<0.05, ** p<0.01, ***p<0.001 vs the indicated group.

## Supplementary Tables

**Supplementary Table 1:** antibodies used for flow cytometry and immunofluorescence.

| **Antibody** | **Fluorochrome** | **Company** | **Reference** |
| --- | --- | --- | --- |
| **CD3** | FITC | BD | 555274 |
| **CD3** | PE | BD | 555275 |
| **CD3** | PercpCy5 | Invitrogen | 4341614 |
| **CD3** | PB | Biolegend | 100214 |
| **CD4** | FITC | BD | 553650 |
| **CD4** | APC-H7 | BD | 560181 |
| **CD4** | PercpCy5 | BD | 553052 |
| **CD5** | FITC | BioLegend | 100605 |
| **CD5** | APC | BioLegend | 100626 |
| **CD8a** | PE | BD | 553032 |
| **CD8a** | PE-Cy7 | eBioscience | 25-0081-81 |
| **CD11b** | PerCP | BioLegend | 101230 |
| **CD11b** | PB | BioLegend | 101223 |
| **CD11b** | AF700 | BioLegend | 101222 |
| **CD11b** | PEDazzle | BioLegend | 101256 |
| **CD16/32** | APC-Cy7 | BioLegend | 101327 |
| **CD19** | PE | eBioscience | 12-0193-82 |
| **CD19** | FITC | Biolegend | 152404 |
| **CD19** | PE-Dazzle594 | Biolegend | 115554 |
| **CD23** | PE | BioLegend | 101608 |
| **CD23** | AF700 | Biolegend | 101631 |
| **CD24** | APC | BioLegend | 101814 |
| **CD25** | BV510 | BioLegend | 101831 |
| **CD38** | PE Cy7 | BioLegend | 102717 |
| **CD38** | PercpCy5 | BioLegend | 102721 |
| **CD45.1** | FITC | eBioscience | 11-0453-82 |
| **CD45.2** | APC | BD | 561875/558702 |
| **CD45.2** | BV510 | Biolegend | 109837 |
| **CD45.2** | BV711 | Biolegend | 109847 |
| **CD45R/B220** | PE | BD | 553089 |
| **CD45R/B220** | PB | Biolgend | 103230 |
| **CD45R/B220** | BUV395 | BD Horizon | 563793 |
| **CD45R/B220** | BV510 | Biolegend | 103247 |
| **CD115** | PE | Biolegend | 135505 |
| **CD115** | APC | eBioscience | 17-1152-82 |
| **CD127 (IL-7Rα)** | PE-Dazzle594 | Biolegend | 135032 |
| **CD138** | AlexaFluor647 | BioLegend | 142525 |
| **CD138** | PE-Dazzle594 | BioLegend | 142527 |
| **CXCR5 (CD185)** | PE | Biolegend | 145513 |
| **CCR6 (CD196)** | PercpCy5 | BioLegend | 129809 |
| **F4/80** | PE | BioLegend | 123110 |
| **IgM** | PE/Cy7 | BioLegend | 406514 |
| **IgM** | AlexaFlour647 | BioLegend | 406525 |
| **IgD** | BV605 | BioLegend | 405727 |
| **Ly6C** | FITC | BD | 553104 |
| **Ly6C** | PE-Cy7 | BD | 560593 |
| **Ly6C** | BV421 | Biolegend | 128031 |
| **Ly6G** | FITC | Biolegend | 127605 |
| **Ly6G** | APC/Cy7 | Biolegend | 127623 |
| **Ly6G** | BUV395 | BD Horizont | 563978 |
| **PNA** | FITC | Vector | FL-1071 |
| **yd TCR** | APC | Biolegend | 17-5711 |

**Supplementary Table 2:**
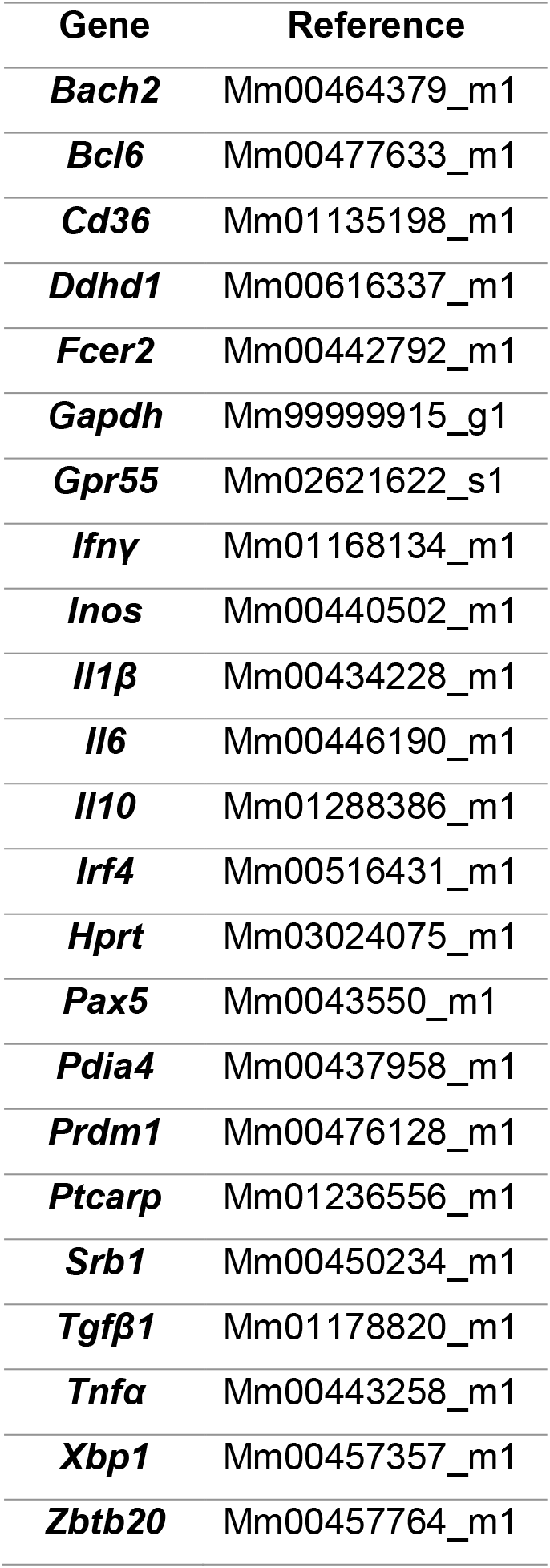
TaqMan Gene Expression Assays used for real time PCR.

**Supplementary Table 3:**
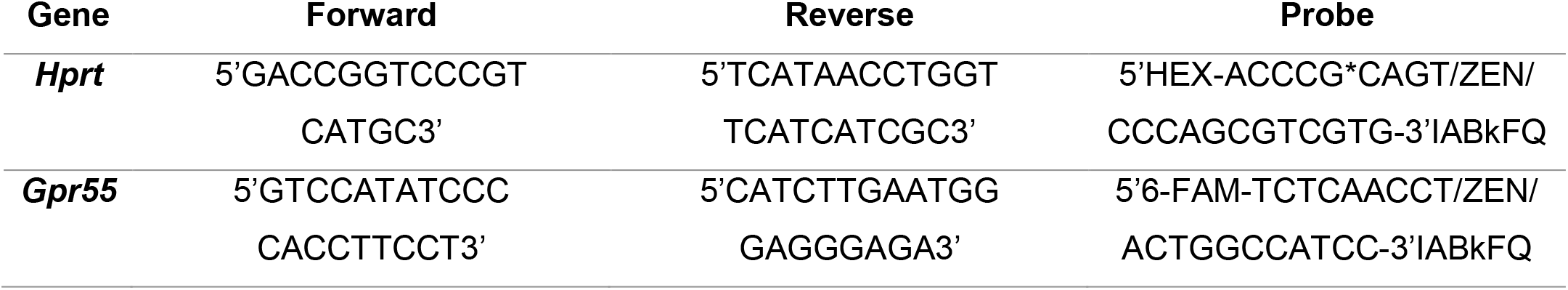
Digital droplet PCR primers and probes (Integrated Data Technologies)

**Supplementary Table 4:** Patient characteristics.

|  | Stable plaque | Unstable/ruptured plaque |
| --- | --- | --- |
| **Number of patients** | 8 | 8 |
| **Age (median)** | 73 | 78 |
| **BMI (median)** | 26.2 | 26.4 |
| **Diabetes mellitus (absolute numbers)** | 2/8 | 1/8 |
| **Sex** | 2 female / 6 male | 3 female / 5 male |
| **Arterial hypertension (absolute numbers)** | 6/8 | 6/8 |
| **Coronary artery disease (absolute numbers)** | 4/4 | 4/4 |
| **Hyperlipidemia (absolute numbers)** | 6/8 | 7/8 |
| **Smoking** | 5/8 | 5/8 |
| **Symptomatic (stroke, TIA, amaurosis fugax; absolute numbers)** | 3/8 | 2/8 |

